# Genome wide inherited modifications of the tomato epigenome by trans-activated bacterial CG methyltransferase

**DOI:** 10.1101/2024.04.17.589930

**Authors:** Bapatla Kesava Pavan Kumar, Sébastien Beaubiat, Chandra Bhan Yadav, Ravit Eshed, Tzahi Arazi, Amir Sherman, Nicolas Bouché

**Author notes:** To whom correspondence should be addressed. Nicolas BOUCHÉ; Amir SHERMAN.

## Abstract

Epigenetic variation is mediated by epigenetic marks such as DNA methylation occurring in all cytosine contexts in plants. CG methylation plays a critical role in silencing transposable elements and regulating gene expression. The establishment of CG methylation occurs via the RNA-directed DNA methylation pathway and CG methylation maintenance relies on METHYLTRANSFERASE1, the homologue of the mammalian DNMT1. Here, we examined the capacity to stably alter the tomato genome methylome by a bacterial CG-specific *M.SssI* methyltransferase expressed through the LhG4/pOP transactivation system. Methylome analysis of *M.SssI* expressing plants revealed that their euchromatic genome regions are specifically hypermethylated in the CG context, and so are most of their genes. However, changes in gene expression were observed only with a set of genes exhibiting a greater susceptibility to CG hypermethylation near their transcription start site. Unlike gene rich genomic regions, our analysis revealed that heterochromatic regions are slightly hypomethylated at CGs only. Notably, some *M.SssI*-induced hypermethylation persisted even without the methylase or transgenes, indicating inheritable epigenetic modification. Collectively our findings suggest that heterologous expression of *M.SssI* can create new inherited epigenetic variations and changes in the methylation profiles on a genome wide scale. This open avenues for the conception of epigenetic recombinant inbred line populations with the potential to unveil agriculturally valuable tomato epialleles.

## INTRODUCTION

Epigenetic variation is mediated by epigenetic marks such as cytosine DNA methylation which occurs in three different contexts CG, CHG, or CHH (H = A, T or C) in plants (Law and Jacobsen, 2010). DNA methylation plays a critical role in silencing Transposable Elements (TEs) and regulating gene expression (Lucibelli et al., 2022). DNA methylation patterns are regulated by various physiological and developmental stimuli, including environmental stresses (Arora et al., 2022). In plants, the establishment of DNA methylation, including at CG sites, occurs via the RNA-directed DNA Methylation (RdDM) pathway, which involves the DOMAINS REARRANGED METHYLTRANSFERASE2 (DRM2) enzyme. Methylation maintenance of CG sites mainly relies on METHYLTRANSFERASE1 (MET1), the plant homologue of the mammalian DNMT1 enzyme. CG methylation occurs in both TEs and genes, leading to the formation of gene body methylation. However, the exact function of gene body methylation is currently unknown. CHG and CHH sites are maintained by methylases like CHROMOMETHYLASE2 (CMT2), CMT3 and DRM2 (Law and Jacobsen, 2010). Out of the three cytosine methylation contexts, the most frequent, heritable, and less influenced by environmental factors is the symmetric methylation of CGs. In tomato, 80% of the CG sites display methylation (Corem et al., 2018), whereas rice exhibits a 40% global methylation rate (Hu et al., 2014), and *Arabidopsis* 24% (Cokus et al., 2008).

Epialleles are alternative epigenetic forms of a specific locus that can potentially influence gene expression and be inherited across generations (Weigel and Colot, 2012). Natural epialleles were identified in plants such as the tomato *COLORLESS NON-RIPENING* (*CNR*) impairing fruit ripening (Manning et al., 2006), *SP11* which is a *B. rapa* epiallele involved in self-incompatibility (Shiba et al., 2006) or the *Lcyc* epiallele involved in flower symmetry of toadflax (Cubas et al., 1999) in addition to several *Arabidopsis* epialleles (Jacobsen and Meyerowitz, 1997; Durand et al., 2012; Agorio et al., 2017). Most studies of epigenetic variation in plants are based on stripping the methylation by chemicals such as 5-Azacytidine or genetic means (*i.e.* mutants) and using the hypomethylated plants as a source of variation to study gene function and isolate new epialleles (Lieberman-Lazarovich et al., 2022). Our comprehension of the mechanisms involved in the creation and maintenance of epialleles was greatly advanced by the creation of epigenetic Recombinant Inbred Lines (epiRILs) which were generated by crossing wild-type *Arabidopsis* accessions with DNA hypomethylated mutants like *met1* or *decreased DNA methylation1* (*ddm1*) (Reinders et al., 2009; Johannes et al., 2009). Still, most of the studies exploring the significance and function of epigenetic modifications have primarily employed *Arabidopsis* as a model plant and relayed on reduction of DNA methylation to generate novel epigenetic variations.

An alternative approach to generating novel plant epialleles is by introducing foreign methylases to induce methylation. This is grounded in the belief that the inherent biological processes will preserve these changes over time. Expression in tobacco of the *E. coli* dam methylase leads to high adenosine methylation at GATC sites and a set of biological phenotypes (van Blokland et al., 1998) demonstrating that a bacterial methylase can methylate plant DNA *in-vivo*. In another work, a foreign methylated DNA could be maintained into tobacco by the plant machinery (Weber et al., 1990), providing evidence that plants can recognize and maintain *de novo* methylated sites. *M.SssI* from the *Mollicutes spiroplasma* species is a bacterial methylase that catalyzes specifically CG methylation (Renbaum et al., 1990). *M.SssI* was shown to be active *in vitro*, associated with Zinc Finger (ZF) proteins (Xu and Bestor, 1997; Chaikind and Ostermeier, 2014), triple-helix-forming oligonucleotides (van der Gun et al., 2010) or catalytically-inactive Cas9 (dCas9) (Lei et al., 2017) and *in vivo* with dCas9 in *E. coli* (Xiong et al., 2018; Ślaska-Kiss et al., 2021), mammalian cells (Xiong et al., 2017), mouse oocytes or embryos (Yamazaki et al., 2017) or with Transcription Activator-Like Effector (TALE) fusion proteins in mouse (Yamazaki et al., 2023). The ability of different *M.SssI* variants fused to a dCas9 to induce methylation in a specific locus was also demonstrated in *Arabidopsis* (Ghoshal et al., 2021), as well as the potential of the newly acquired methylation to be inherited. The same *M.SssI* variant fused to an artificial ZF domain induces methylation in specific and nonspecific modes that were also inherited by the next generations (Liu et al., 2021). However, till now, utilizing native *M.SssI* to induce genome scale CG methylation in plants was not reported. In this study, we overcame difficulties to express native *M.SssI* in tomato using a two-component transcription activation system (Moore et al., 1998). Analysis of the methylome of the trans-activated plants expressing *M.SssI* revealed that the expression of *M.SssI* devoid of fusion proteins has significant repercussions on the overall methylation homeostasis of tomato even when the transgenes were segregated away in the following generations.

## RESULTS

### Ectopic expression of a bacterial DNA methylase in tomato

*M.SssI* is a bacterial CG methyltransferase with specific codon usage (Renbaum *et al*., 1990). To constitutively express *M.SssI in planta*, we optimized its codon usage and added a nuclear localization signal in-frame at the 3’-end. The potato *IV2* intron (Eckes et al., 1986) was introduced into the plant-adapted *M.SssI* coding sequence to prevent bacteria from expressing the active enzyme (Sup Figure 1) facilitating its cloning into a binary plasmid. The disarmed plant-adapted *M.SssI* (here after named *disM.SssI*) was cloned in front of a double *CaMV 35S* promoter followed by a TMV omega leader sequence to constitutively express it and assist its translation, respectively (Methods). To test whether the *disM.SssI* enzyme is active *in planta*, we transiently expressed it in *Nicotiana benthamiana* leaves and quantified the global cytosine methylation levels of their genomic DNA two days after infiltration (2 dpi). The *pART27_2×35S_Omega_disM.SssI* infiltrated leaves showed a significant increase in global 5-methyl cytosine compared to the empty vector infiltrated leaves (Sup Figure 2). This result suggests that *disM.SssI* was active *in planta*. Transformation of *Arabidopsis* and tomato plants with *Agrobacterium* carrying the *pART27_2×35S_Omega_disM.SssI* binary plasmid, repeatedly failed to recover transgenic plants, suggesting that constitutive expression of *disM.SssI* might be lethal to plants.

**Figure 1.**
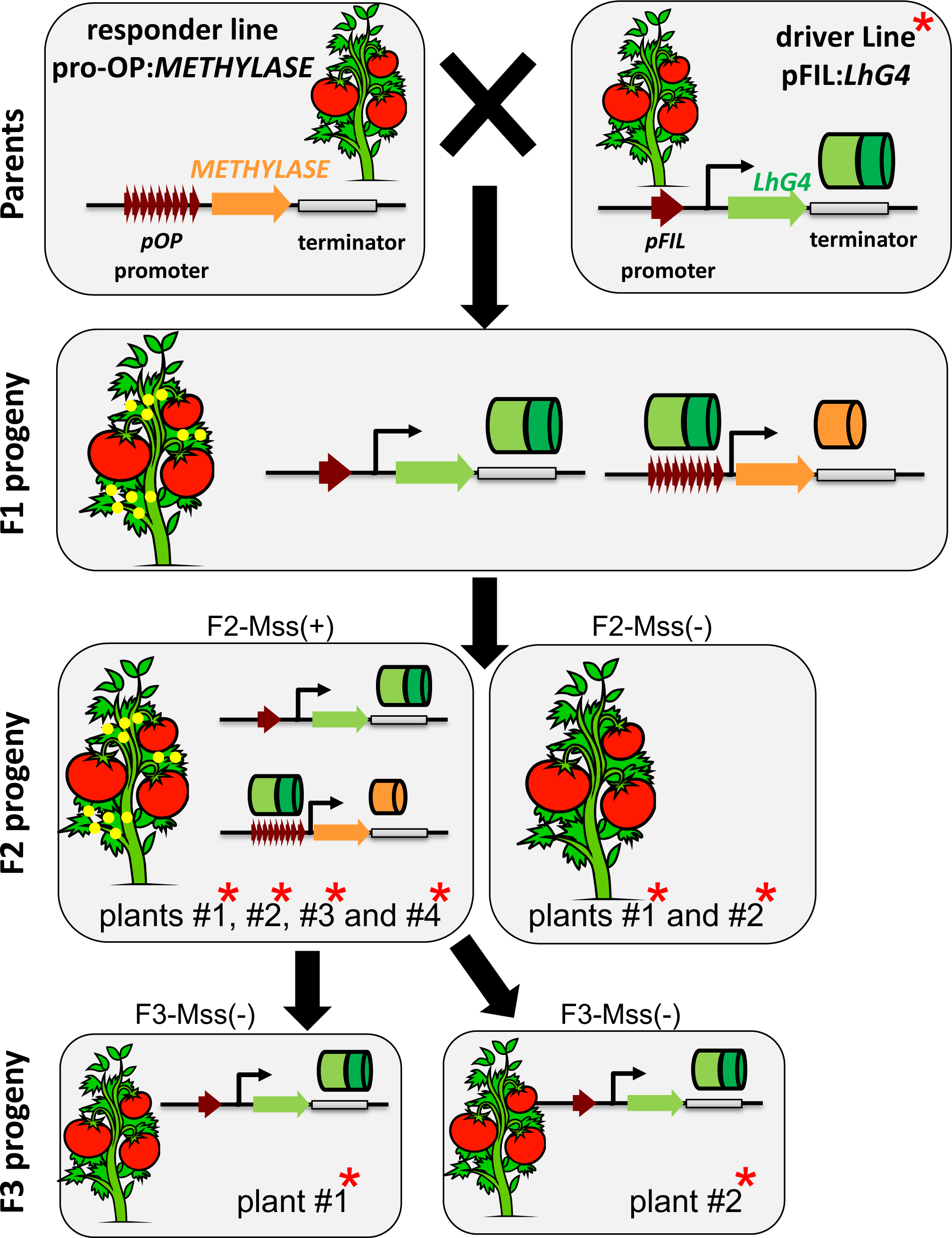
Schematic representation of the different plants produced in this study. Yellow sphere, expression of the *M.SssI* enzyme. Red asterisk, plant methylomes sequenced in this study.

**Figure 2.**
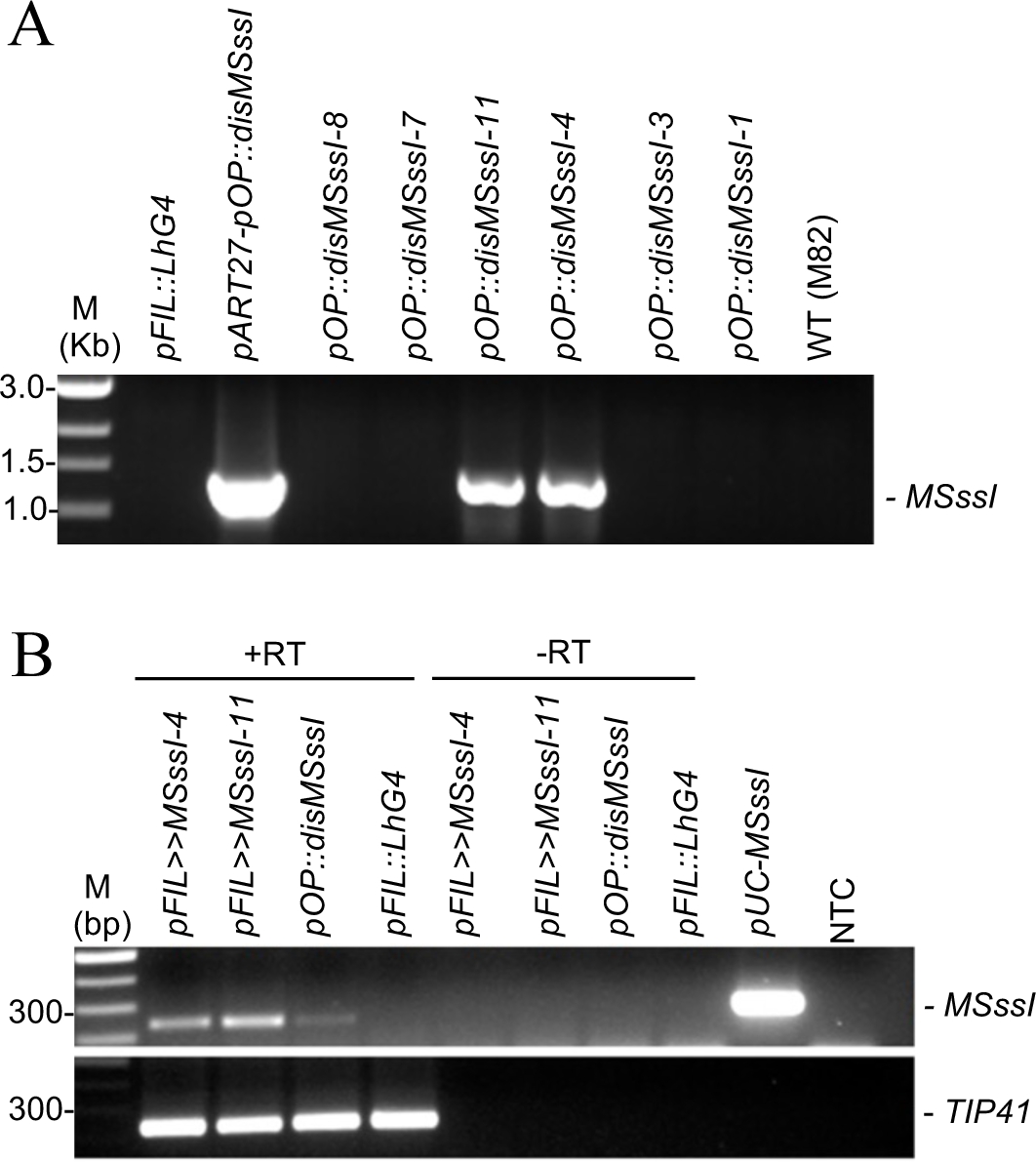
Transactivation of *pOP::disMSssI* results in its expression in tomato. (A) Genotyping of transgenic tomato plants (F1 generation) by PCR to test for the presence of the *pOP::disM.SssI* transgene. The plasmid *pART27-pOP::disM.SssI* was used as a positive control and the plasmid *pFIL::LhG4* construct as a negative control. Wild-type DNA (WT) correspond to DNA extracted from M82 cultivar leaves. M, DNA marker. (B) Expression of *disM.SssI* analysed by RT-PCR in two different F1 lines carrying both transgenes (*pFIL>>pOPdisM.SssI-4 and 11)*. The absence of Reverse Transcriptase (-RT) was used for negative control and the *pUC::disM.SssI* plasmid DNA as a positive control. The *TIP41* gene was used as an equal loading control. M, DNA marker. NTC, No Template Control.

To facilitate the expression of *M.SssI* in tomato, we utilized the LhG4/pOP transactivation system, which separates the transformation and transgene expression steps (Moore et al., 1998). Two independent *pOP::disM.SssI* transgenic M82 responder lines, were obtained by transformation and regeneration (Methods). The *pOP* promoter is normally inactive and is *trans*-activated only in the presence of its artificial *pOP* activator *LhG4* (Figure 1). To induce the expression of *disM.SssI*, the *pOP::disM.SssI* responder lines carrying the construct (Figure 2A) were crossed with a homozygous *pFIL::LhG4* driver line expressing *LhG4* under the *FILAMENTOUS FLOWER* (*FIL*) promoter that was described as primordia and leaf specific (Lifschitz et al., 2006). The F1 transactivated progenies (*pFIL::LhG4 >> pOP::disM.SssI*) germinated normally and overexpressed the *disM.SssI* transgene (Figure 2B). Although their cotyledons were not different from that of wild-type plants, some F1s developed severely distorted leaves consistent with the *pFIL* expression domain (sup Figure 3) and reminiscent of the tomato *wiry* phenotype (Yifhar et al., 2012). All *pFIL>>disM.SssI* plants were fertile and further analyses were done on their F2 progeny (Figure 1).

**Figure 3.**
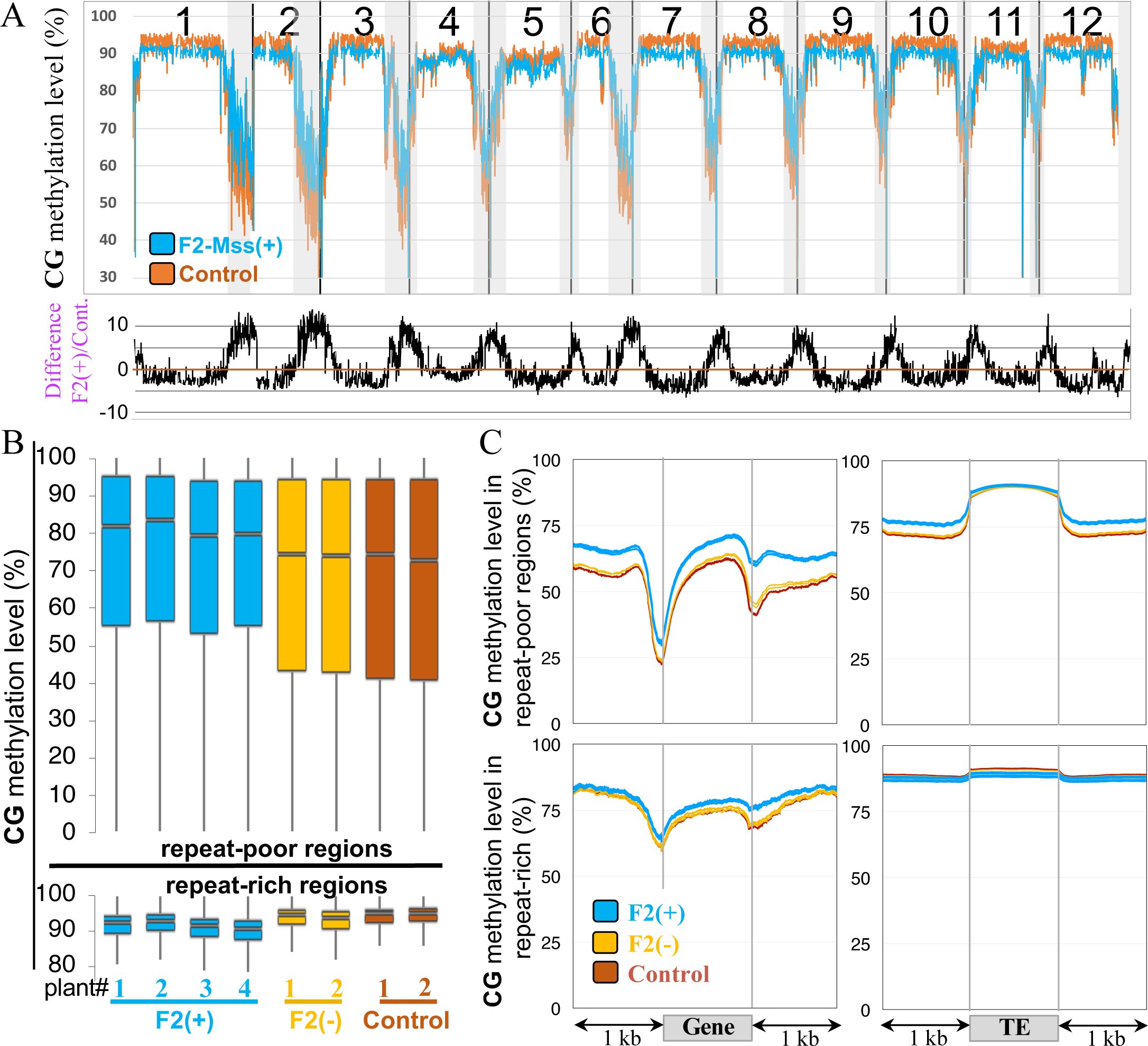
CG methylation in tomato F2 plants expressing or not *M.SssI*. (A) Methylation across the 12 chromosomes of tomato determined for non-overlapping 200 kb-bins that cover the entire genome. The methylation levels correspond to the proportions of methylated cytosines relative to the total number of cytosines calculated by aggregating the outcomes from all F2-Mss(+) or control plants. The regions enriched for genes are in grey. (B) Box plots showing mean methylation content of the F2-Mss(+), F2-Mss(-) and control lines. The SL2.50 version of the tomato genome assembly (Tomato Genome Consortium, 2012) was segmented in 1kb windows; methylation levels correspond to the proportions of methylated cytosines relative to the total number of cytosines. Only cytosines covered by at least five reads were considered and only bins containing at least 10 valid cytosines were considered. The repeat-rich and repeat-poor regions were defined as previously described (Jouffroy et al., 2016). Control: *pFIL::LhG4* driver lines. The correlation network diagram constructed using Spearman’s correlation coefficients to illustrate the relationships among the CG methylation bins depicted in the boxplots is shown in Sup Figure 6. (C) Metaprofiles of CG methylation for genes and Transposable Elements (TEs). TEs were annotated with REPET (Flutre et al., 2011).

### Expressing *M.SssI* increases the CG methylation of tomato genes

To study the impact of the expression of *M.SssI* on tomato genome methylation, the methylomes of plants expressing the transgene were sequenced. Genomic DNA was extracted from leaves of F2 progenies in which the transgenes were segregating to sequence the methylomes of four individual F2 plants carrying both the *pFIL:LhG4* and the *pOP::disM.SssI* transgenes (hereafter named F2-Mss(+) plants), as well as two other F2 sibling plants containing no transgenes (hereafter named F2-Mss(-) plants). The potential significance of methylome variation throughout transformation, across various generations, genotypes, and individuals cannot be understated, hence, it is crucial to reduce these variances through the utilization of a suitable control. We therefore conducted methylome sequencing for two individual *pFIL::LhG4* driver plants cultivated alongside the F2 plants, along with an additional two independent *pFIL::LhG4* driver plants grown at a separate instance. All the following analyses were carried out using these four control plants (hereafter named control driver line plants). The alignment of the reads with the sequences of the *pFIL::LhG4* or the *pOP::disM.SssI* transgenes reconfirmed that all plants belong to the different genotypes analysed (Sup Figure 4). The levels of methylation per cytosine were determined for all methylation contexts (CG, CHG and CHH). On the chromosomal scale, DNA methylation was assessed by calculating methylation levels within 200 kb-windows that covered the entire genome. The results were then graphically represented by mapping them onto the 12 tomato chromosomes (Figure 3A). Within the gene-containing regions of every chromosome, CG methylation was globally increased for the F2-Mss(+) plants expressing the bacterial methylase, compared to control driver line plants or F2-Mss(-) (Figure 3A, grey areas and Sup Figure 5A). In centromeric and pericentromeric regions enriched for TEs, the level of CG methylation exhibited a minor reduction for F2-Mss(+) plants but not for F2-Mss(-) plants (Figure 3A, white areas and Sup Figure 5). Methylation levels at chromosome scales were similar for the CHG contexts between F2-Mss(-), F2-Mss(+) and control driver line plants (Sup Figure 5B and C). We also note that all F2 plants seem to be slightly hypermethylated in the CHH context compared to driver line plants (Sup Figures 5D and E). These findings were confirmed when the average methylation levels were calculated within 1 kb-segments dividing the genome. Regions characterized by a high gene density and a low number of TEs (i.e. repeat-poor regions as described in Jouffroy et al., 2016) were hypermethylated in the CG context for the four individual F2-Mss(+) plants, contrarily to the F2-Mss(-) or to the control driver line plants (Figure 3B, *repeat-poor regions*; Sup Figure 6). On the opposite, regions containing a low number of genes and densely populated by TEs (i.e. repeat-rich regions as described in Jouffroy et al., 2016) were hypomethylated in the CG contexts only in F2-Mss(+) plants (Figure 3B, *repeat-rich regions*; Sup Figure 6). Therefore, methylome sequencings of leaves revealed a global increase of CG methylation in genic (repeat-poor) regions and a decrease of CG methylation in heterochromatic (repeat-rich) regions. Consequently, the introduction of the bacterial *M.SssI* enzyme into tomato resulted in a widespread disruption of the CG methylation balance.

**Figure 4.**
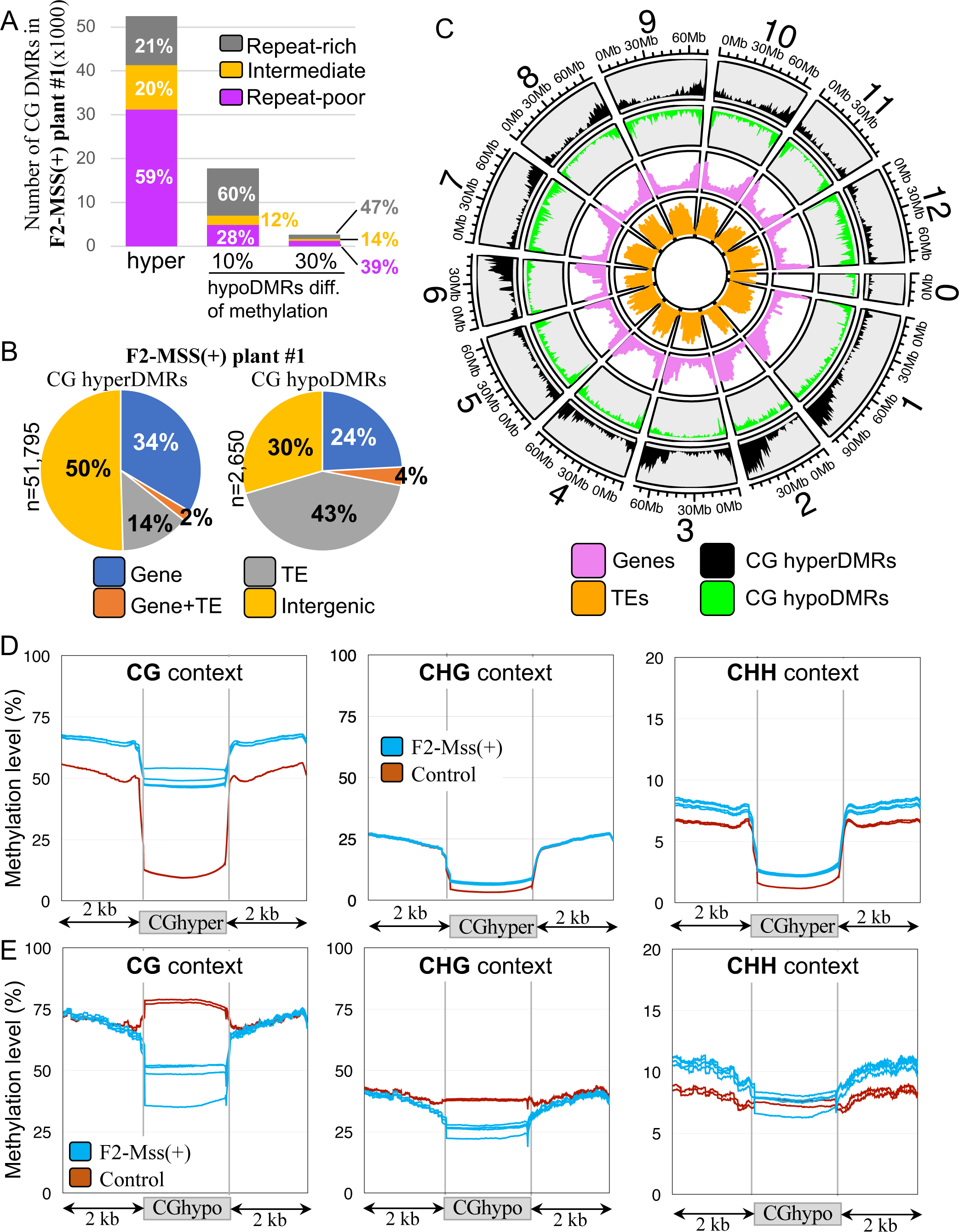
Nature and localization of the Differentially Methylated Regions (DMRs) identified between the F2-Mss(+) plant#1 are the driver line control plants for the CG methylation context. (A) Hypermethylated (n=51,795) CG DMRs identified between the F2-Mss(+) plant#1 and the driver line control plants when a 30% absolute difference of methylation was applied and hypomethylated DMRs identified using a 30% (n=2,650) or a 10% (n=17,508) absolute difference of methylation. All other three F2-Mss(+) plants show similar numbers (Sup Table 2 and Sup Figure 9). The repeat-rich, repeat-intermediate and repeat-poor regions, based on the repeat densities, were defined as previously described (Jouffroy et al., 2016). (B) Nature of the CG hypomethylated and hypermethylated DMRs identified between the F2-Mss(+) plant#1 and the driver line control plants. “CDS+TE” are DMRs overlapping with both CDSs and transposons, “CDS”, DMRs overlapping with CDSs, and “TE” DMRs overlapping with Transposable Elements (TEs). All other DMRs were classified as “Intergenic”. The nature of the CG DMRs identified between all F2-Mss(+) plants and the driver line control plants is shown in Sup Figure 10. (C) Densities of CG hypomethylated (*green*) and hypermethylated (*black*) DMRs identified between the F2-Mss(+) plant#1 and the driver line controls along the 12 tomato chromosomes. The density of genes and TEs are shown in pink and orange, respectively. (D) Metaprofiles of methylation in the three methylation contexts (CG, CHG and CHH) for the CG hypermethylated and hypomethylated identified between the F2-Mss(+) plant#1 and the driver line control plants.

**Figure 5.**
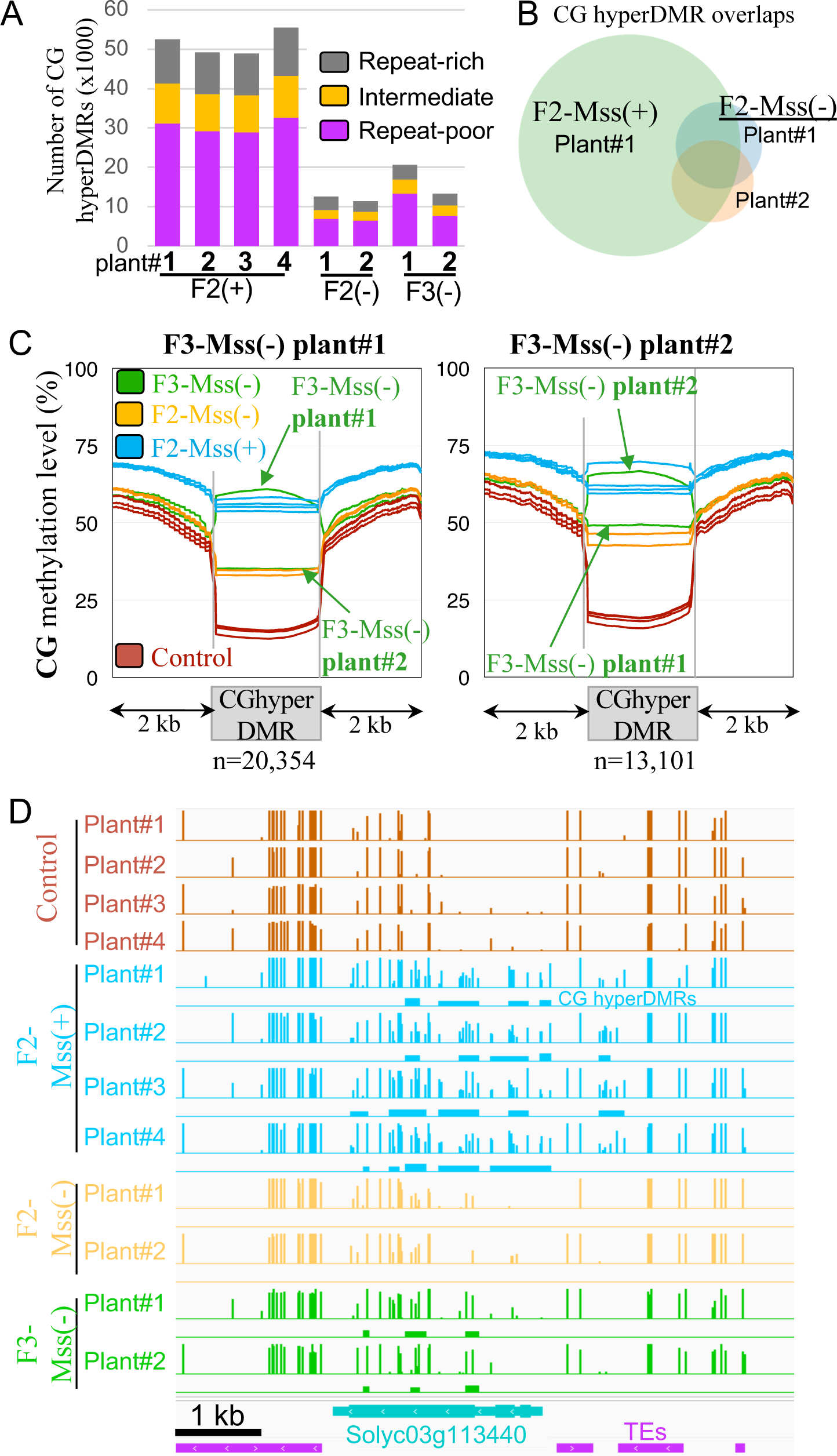
CG hypermethylated DMRs induced by *M.SssI* are transmitted between tomato generations. (A) Hypermethylated CG DMRs detected in F2 and F3 plants compared to the driver line control plants. The repeat-rich, repeat-intermediate, and repeat-poor regions, based on the repeat densities, were defined as previously described (Jouffroy et al., 2016), using the SL2.50 version of the genome assembly. The numbers of DMRs for all plants are shown in Sup Figure 9. (B) Overlap of hypermethylated CG DMRs between F2-Mss(+) plant#1, F2-Mss(-) plant#1 and F2-Mss(-) plant#2. (C) Methylation levels of plant#1 and plant#2 F3-Mss(-) CG hyperDMRs. The average methylation levels were determined by dividing the DMRs into 100-bp bins. Methylation levels in regions located 2 kb upstream and 2 kb downstream the DMRs are shown. (D) Example of genome view of CG methylation patterns in F2 and F3. CG hyperDMRs between the control driver line plants and the different genotypes are shown as colored rectangles below the methylation track.

**Figure 6.**
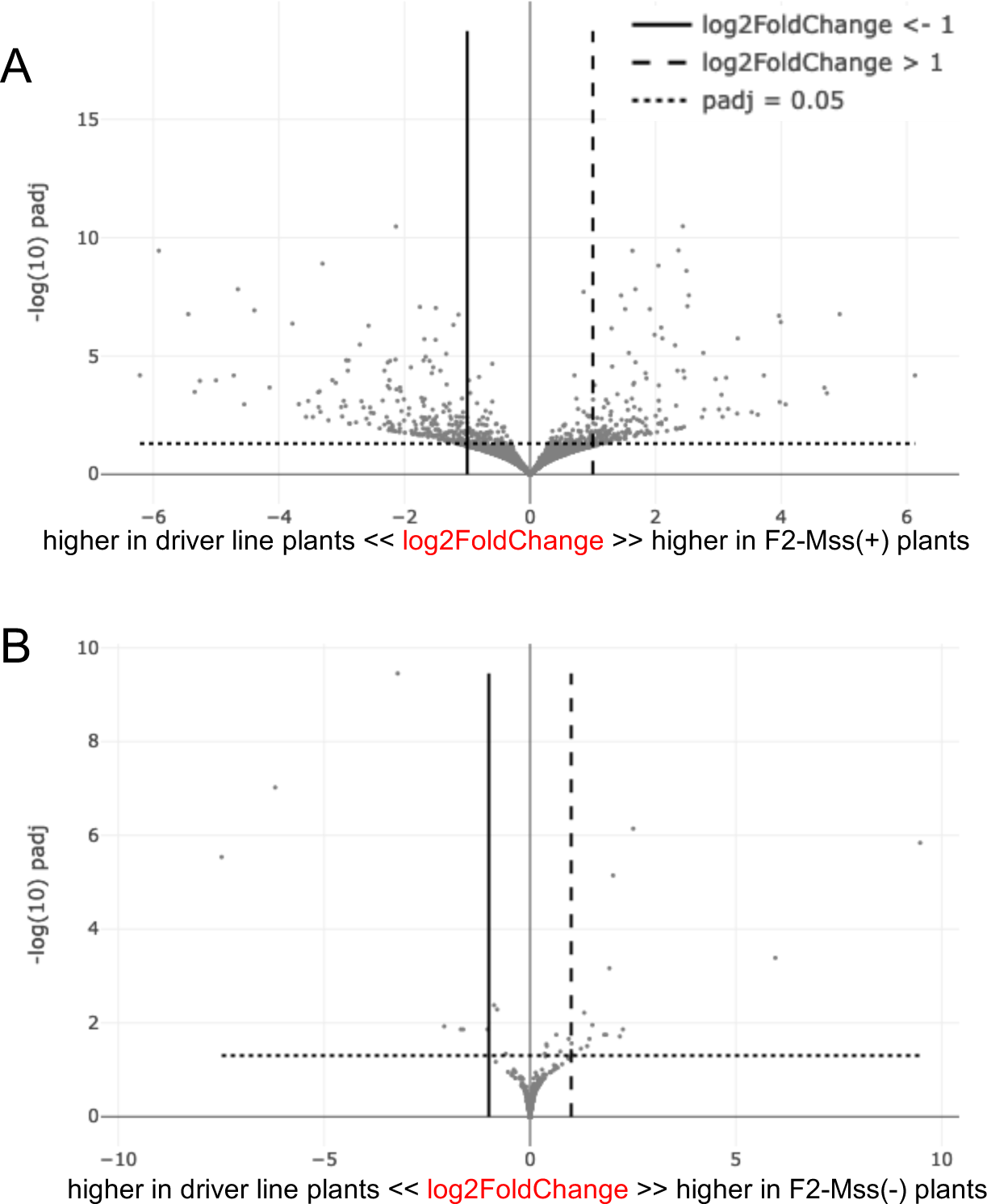
Differences of gene expression in F2-Mss(+) plants (A) and F2-Mss(-) plants (B) compared to the control lines, represented by volcano plots of −log(10) p-value against log(2) fold changes for Differentially Expressed Genes (DEGs).

On average, genes were hypermethylated in the CG context in F2-Mss(+) plants compared to F2-Mss(-) or driver line control plants (Figure 3C). The differences in methylation were more pronounced in the 3’-end part of the genes and were observed for genes localized in both repeat-poor or -rich regions (Figure 3C). No differences were observed in the CHG context (Sup Figure 7). In the CHH context, genes localized in repeat-rich regions are more methylated in both F2-Mss(+) and F2-Mss(-) plants compared to controls (Sup Figure 7). On the other hand, the methylation of TEs was similar between F2-Mss(+) and F2-Mss(-) plants, in all symmetric methylation contexts (Sup Figure 8). Again, the CHH context is an exception and TEs have increased methylation compared to control driver lines, whether they are localized in heterochromatic or euchromatic regions (Sup Figure 8). Altogether, the data show a specific increase of CG methylation in genes for plants expressing the bacterial *M.SssI* methylase.

**Figure 7.**
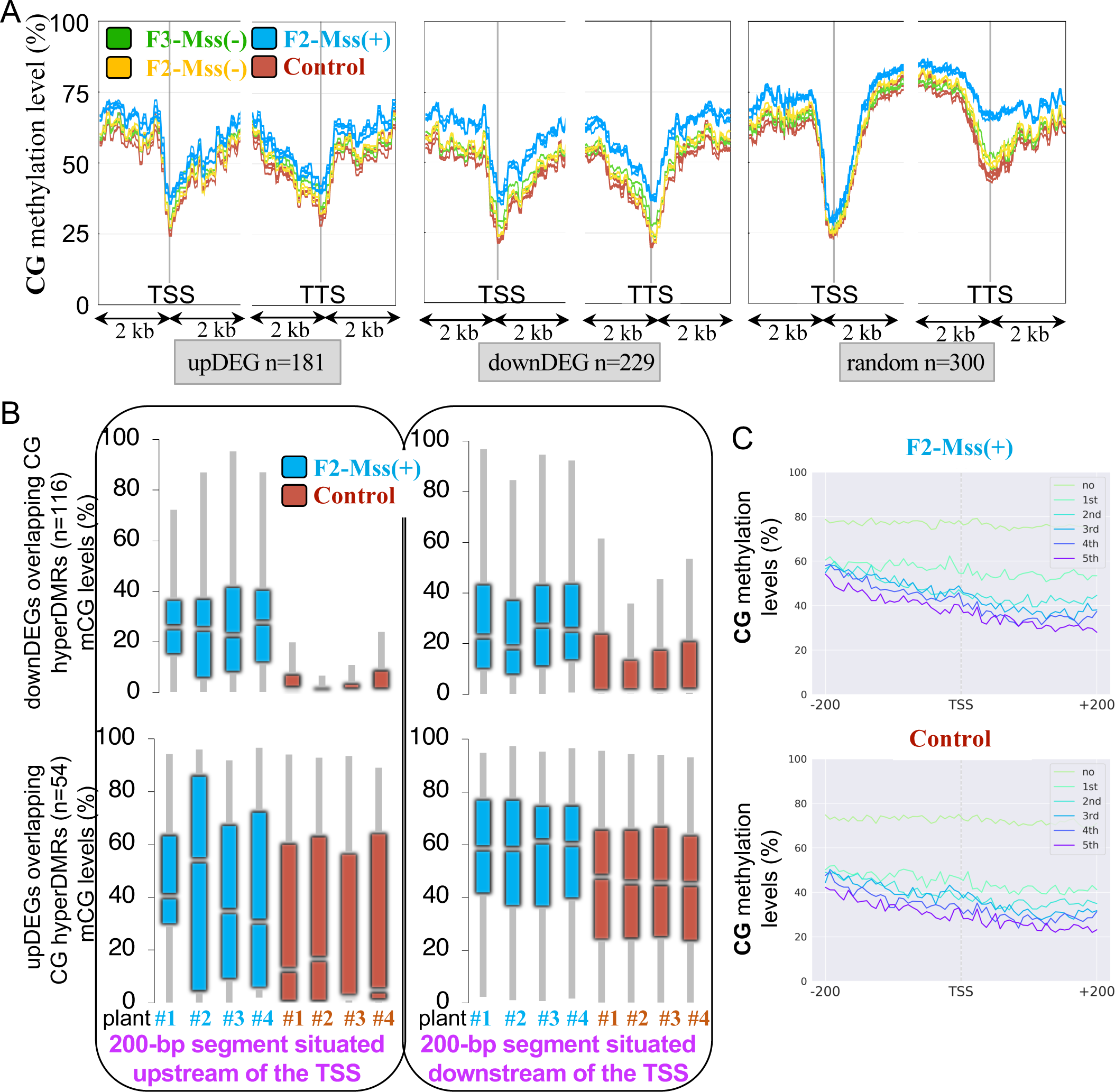
Methylation levels of Differentially Expressed Genes (DEGs) that are up- or downregulated (> ou < 1 log2FC) in F2/F3 plants expressing *M.SssI* (*Mss(+)*) or not (*Mss(-)*) versus the control driver line plants. The average methylation levels of the DEG was calculated by dividing the DEG region into 100-bp bins. Regions located 2-kb upstream and 2-kb downstream the DEGs are shown. Random: set of 300 genes selected randomly. (B) Box plots showing the mean methylation near the Transcription Start Site (TSS) of upregulated DEGs (upDEGs) overlapping with CG hyperDMRs. Only genes with TSS overlapping (+/- 500bp) with CG hyperDMRs were considered. Only genes whose TSS overlap with CG hypermethylated regions within a range of +/- 500-bp were considered. Methylation levels were calculated in regions upstream (−200 bp) and downstream (+200 bp) of F2-Mss(+) and control line TSSs as the proportions of methylated cytosines over the total number of cytosines. Only cytosines covered by at least five reads were considered and only bins containing at least 5 valid cytosines were kept. (C) Average methylation level profiling according to different expression groups around the TSS (+/- 200 bp) of F2-Mss(+) and control lines. Genes are grouped as non-expressed genes and five quantiles of expressed genes according to the gene expression level groups from low to high; the first quintile is the lowest, and the fifth is the highest.

### *M.SssI* targets unmethylated genic regions and accessible chromatin

The regions that were significantly differentially methylated (Differentially Methylated Regions, DMRs) between F2-Mss(+) or F2-Mss(-) plants and the driver line controls were identified. Compared to the driver line control plants, the F2-Mss(+) plants contained the highest number of DMRs with a vast majority of hypermethylated DMRs (hyperDMRs) in the CG context (Plant#1: n=51,795; Plant#2: n=48,467; Plant#3: n=48,254; Plant#4: n=54,758), consistent with *M.SssI* being active in these plants (see Figure 4A for the example of F2-Mss(+) plant#1 and Sup Figure 9 for all plants). CG hyperDMRs mainly overlapped with intergenic regions (50% of the CG hyperDMRs in F2-Mss(+) plant#1, Figure 4B and Sup Figure 10) and genes (34% of the CG hyperDMRs in F2-Mss(+) plant#1, Figure 4B and Sup Figure 10). In agreement with this last observation, 59% of the CG hyperDMRs of F2-Mss(+) plant#1 (n=31,153) are found within repeat-poor regions enriched for genes, containing low amounts of repeats (Figure 4A, *Repeat-poor regions*) and localized near chromosome arms (Figure 4C). Moreover, ∼34% of the tomato genes overlap with at least one CG hyperDMRs in the F2-Mss(+) plant#1 with similar results obtained for the three other plants analysed (32% for F2-Mss(+) plant#2, 33% for F2-Mss(+) plant#3 and 35% for F2-Mss(+) plant#4). The number and localization of the DMRs identified between the other F2-Mss(+) plants (#2, #3 and #4) and the controls are similar to what was found for F2-Mss(+) plant#1 (Sup Table 2 and Sup Figure 9). Altogether, our findings demonstrate that a significant portion of tomato genes undergo CG methylation when the corresponding plants ectopically express the bacterial *M.SssI* methylase.

Most of the CG hyperDMRs (96% n=49,762) found between F2-Mss(+) plant#1 and the controls are not overlapping with hypermethylated or hypomethylated DMRs (hypoDMRs) in other cytosine contexts, namely CHG or CHH. Hence, most regions that experience an increase in CG methylation upon methylase expression do not undergo significant alterations in their methylation patterns for other types of DNA methylation. This was confirmed when the metaprofiles of methylation were drawn for the CG hyperDMRs of F2-Mss(+) plants (Figure 4D, Sup Figures 11 and 12). Indeed, CG hyperDMRs observed in F2-Mss(+) plants are indicative of regions where the control driver line plants exhibit minimal levels of basal methylation. On average, these regions are methylated at 10% for CGs, 3.7% for CHGs and 1.5% for CHHs in the control lines, reaching about 50% of CGs methylated in individual F2-Mss(+) plants with a significant (t-test p-value<0.005) increase of 372% compared to the control (Figure 4D, Sup Figures 11 and 12). Both CHG and CHH sites gain methylation to a much lesser extent with a significant (t-test p-value<0.005) increase of 90% for CHGs and 60% for CHHs (Figure 4D, Sup Figures 11 and 12). Altogether, the data indicate that the bacterial methylase targets preferentially euchromatic regions that were almost unmethylated in the wild-type tomato genome. In agreement with this hypothesis, we found that 44% (n=23,014 for F2-Mss(+) plant#1) of the CG hyperDMRs overlap with accessible chromatin regions revealed genome-wide using ATAC-seq (Assay for Transposase-Accessible Chromatin using sequencing) data available for tomato meristem-enriched tissues (Hendelman et al., 2021). Accessible chromatin regions correspond to 10% of the total genome sequence length (Hendelman et al., 2021).

There are almost 20 times less CG hypoDMRs (n=2,650 for F2-Mss(+) plant#1) localized more predominantly in heterochromatin (Figure 4A) and mostly matching TEs (Figure 4B). In addition, CG hypoDMRs also seem to be slightly hypomethylated in the CHG but not in the CHH context (Figure 4E). When the threshold of CG methylation difference was lowered from 30% to 10%, we observed a significant increase in the number of detected CG hypoDMRs (Plant#1: n=17,508; Plant#2: n=14,924; Plant#3: n=26,189; Plant#4: n=31,441). Between 56% and 68% of those hypoDMRs were found in repeat-rich regions (Figure 4A), overlapping TEs. The results show a shift in CG methylation from heterochromatin to euchromatic regions, in agreement with chromosome scale observations (Figure 3A).

### CG hyperDMRs are transmitted between tomato generations

To ascertain whether the inheritance of CG hyperDMRs is possible across generations, we examined the methylomes of two F2-Mss(-) non transgenic plants, in which all transgenes from the F1 parent segregated away (Sup Figure 4), and that were cultivated and subjected to sequencing alongside the F2-Mss(+) plants and inherited from the same F1. On average, the CG methylation of both genes and TEs was very similar genome-wide between the F2-Mss(-) non transgenic plants and the driver line controls (Sup Figures 7 and 8). DMRs were identified, revealing that F2-Mss(-) plants still contain a high number of CG hyperDMRs (n=12,410 for F2-Mss(-) plant#1 and n=11,177 for F2-Mss(-) plant#2; Figure 5A, Sup Figure 9 and Sup Table 2). By contrast, the number of CG hypoDMRs was much more limited (n=745 for F2-Mss(-) plant#1 and n=991 for F2-Mss(-) plant#2; Sup Figure 9 and Sup Table 2). As observed for the F2-Mss(+) plants, the CG hyperDMRs of F2-Mss(-) plants are mostly located within regions enriched for genes (n=6,867 for F2-Mss(-) plant#1 and n=4,452 for F2- Mss(-) plant#2; Figure 5A). Furthermore, the CG hyperDMRs identified in F2-Mss(-) plants significantly coincide with the CG hyperDMRs present in their F2-Mss(+) counterparts. For instance, 77% of the CG hyperDMRs found in F2-Mss(-) plant#1 and 70% of the CG hyperDMRs found in F2-Mss(-) plant#2 overlap with CG hyperDMRs of F2-Mss(+) plant#1 (Figure 5B). 44% of the CG hyperDMRs are shared among the two F2-Mss(+) plants analysed (Figure 5B). This indicates that the vast majority of CG hyperDMRs detected in F2- Mss(-) non transgenic plants compared to the driver line controls are shared with their F2- Mss(+) transgenic sibling plants, implying that they were likely inherited from the F1 parent.

To track the potential transfer of CG DMRs across successive generations, F2-Mss(+) plant#3 and plant#4 were selfed and the F3 offspring was genotyped for the presence of the *pFIL:LhG4* and the *pOP:disM.SssI* transgenes. The F3 generation showed a classical Mendelian pattern of segregation for both transgenes, indicating that they existed in a heterozygous state in the preceding F2 parents. The methylomes of two F3 plants carrying solely the *pFIL:LhG4* transgene were sequenced and compared to the control driver line plants which are composed of a *pFIL:LhG4* set of plants including two plants grown together with these F3s and two other plants grown independently with F2s, as stated above. In both F3-Mss(-) plants, the CG hyperDMRs exhibited the highest count of DMRs (n=20,354 for the F3-Mss(-) plant#1, a descendant of F2-Mss(+) plant#3 and n=13,101 for the F3-Mss(-) plant#2, a descendant of F2-Mss(+) plant#4; Sup Table S2). Like for F2s, F3-Mss(-) CG hyperDMRs are localized mostly in genic regions (65% of the CG hyperDMRs for F3-Mss(-) plant#1 and 59% for F3-Mss(-) plant#2 are localized in regions containing low amounts of repeats; Figure 5A, Sup Figure 9 and Sup Table 2). To better understand what the identified DMRs in the F3 correspond to, their methylation metaprofiles have been generated. The metaprofiles of CG hyperDMRs of F2-Mss(+) plants show that these regions are also methylated in both their F2-Mss(-) siblings and F3-Mss(-) progenies, but almost unmethylated in the control driver lines. No changes were detected in the two other CHG and CHH methylation contexts (Sup Figures 11 and 12). CG hyperDMRs identified in F3-Mss(-) plants are at levels of methylation identical to the one of their F2-Mss(+) parents (Figure 5C; Sup Figure 13). Accordingly, most of the CG hyperDMRs of F3-Mss(-) plants are shared with their F2-Mss(+) parents (Figure 5D). Indeed, 76% of the CG hyperDMRs of F3-Mss(-) plant#1 and 84% of F3-Mss(-) plant#2 overlap with CG hyperDMRs found in their corresponding F2-Mss(+) parents. These inherited DMRs account for 32% of the overall number of CG hyperDMRs found in F2-Mss(+) plant#3 and 20% of the CG hyperDMRs found in F2-Mss(+) plant#4.

Thus, both F2 and F3 plants lacking the *M.SssI* gene but descended from *pFIL>>disMSssI* plants that express the transgenes, still carry CG hyperDMRs exhibiting retained levels of methylation when compared to their parent plants. This implies the inheritance of methylation patterns.

In agreement with the CHH methylation levels increase monitored in both F2-Mss(+) and F2-Mss(-) (Sup Figures 5D and 5E, Sup Figures 6 and 7), many CHH hyperDMRs were identified between F2-Mss plants and control driver lines (Sup Figure 9). Most of these DMRs overlap with TEs (for instance, 69.7% of the CHH hyperDMRs overlap with TEs in F2-Mss(+) plant #1) found within repeat-poor regions (Sup Table 2). Nonetheless, F3-Mss(-) plants did not exhibit CHH hyperDMRs, suggesting a substantial divergence in the presence of CHH hyperDMRs between Mss(-) plants from distinct generations (F2 and F3). Moreover, CHH hyperDMRs are not overlapping with DMRs found for other methylation contexts (<2% of the CHH hyperDMRs in F2-Mss(+) plant#1 overlap with other CG or CHG DMRs). Thus, hypermethylated CHH regions are largely independent of other methylation contexts and are specific of F2s. This implies that CHH hyperDMRs are likely independent of the *M.SssI* activity.

### Changes in CG methylation patterns correlate with limited effects on expression

To explore whether the accumulation of CG methylation within genes impacts their expression, we performed an RNAseq analysis of F2 plants. RNAs were extracted from leaves of three F2-Mss(+) plants, three F2-Mss(-) plants and three driver line plants grown together (Sup Table 3). Reproducibility between biological replicates (a single replicate corresponds to an individual plant and is created by combining bulk samples of leaves) was confirmed by performing a Principal Component Analysis (PCA) to visualize the differences (Sup Figure 14). Among the genes that were significantly differently regulated (adjusted p-value <0.05 and log2FoldChange <-1 or > 1) between F2-Mss(+) and control driver line plants, 56% (n=229) were downregulated (down Differentially Expressed Genes, downDEGs; Sup Table 4; Figure 6A) and 44% (n=181) were upregulated (up Differentially Expressed Genes, upDEGs; Sup Table 5; Figure 6A). Altogether, up- and downDEGs in F2-Mss(+) plants represent only 1.2% of the tomato genes. This implies that CG hypermethylation changes occurring within or near genes in these plants have limited effects on global tomato gene expression. 48% to 51% of the downDEGs (n=114 for F2-Mss(+) plant#1, n=117 for plant#2, n=109 for plant #3 and n=117 for plant #4) overlap with CG hyperDMRs. By contrast, these numbers drop to 31 to 38% for the upDEGs (n=69 for F2-Mss(+) plant#1, n=65 for plant#2, n=57 for plant#3 and n=66 for plant#4) which is comparable to the average numbers of genes overlapping with at least one CG hyperDMR genome-wide in F2-Mss(+) plants (∼35%). Only a maximum of 12% of the upDEG and 15% of the downDEGs overlap with CHG or CHH DMRs. By contrast, expression analyses of the three F2-Mss(-) plants revealed very few changes compared to control driver lines with only 16 upDEGs and 7 downDEGs (Figure 6B). Thus, downregulated genes in F2-Mss(+) plants appear to exhibit a higher susceptibility to CG hypermethylation compared to upregulated genes.

To further test this hypothesis, we analyzed the metaprofiles of DEGs for DNA methylation. The methylation profiles of downDEGs differ from those of upDEGs, or genes chosen randomly, mainly around the Transcription Start Site (TSS) and only for CG methylation (Figure 7A and Sup Figure 15). DEGs with CG hyperDMRs localised around their TSS (within a TSS distance of +/- 500-bp) in at least one of the F2-Mss(+) plants were further analysed (n=116 for downDEGs and n=54 for upDEGs). Our findings revealed that the 200 bp region located upstream of the TSSs of downDEGs exhibited nearly negligible CG methylation in control lines, in contrast to F2-Mss(+) plants (Figure 7B). The regions localised 200-bp downstream of the TSS of downDEGs were slightly more methylated (Figure 7B). In contrast, the methylation patterns surrounding the TSSs of upDEGs were distinct, showing significantly elevated average levels of CG methylation preceding and following the TSS in both F2-Mss(+) and control plants (Figure 7B). This suggests a correlation between the presence of CG methylation within the region adjacent to the TSS and the transcriptional decrease of the associated genes in F2-Mss(+) plants. Afterwards, the genes were categorized into two groups: non-expressed genes and expressed genes divided into five quantiles based on their sgene expression level, ranging from low to high. The lowest expression level corresponds to the first quintile, while the highest expression level corresponds to the fifth quintile. Metaprofiles were generated for the various gene categories within windows that cover a range of +/- 200 bp around the TSS (Figure 7C). A significant increase in methylation was only detected in the CG context, particularly in genes characterized by low expression levels (genes in the first quintile; Figure 7C and Sup Figure 16). Hence, the genes expressed at lower levels in the wild-type exhibit a higher sensitivity to CG hypermethylation around their TSS.

TEs found in the vicinity of genes can potentially interfere with gene expression (Baduel and Colot, 2021). However, the results reveal that the proportion of TEs overlapping with DEG (for upDEGs, n=75 which represent 41.4% of all DEGs and for downDEGs n=111 which represent 48,5% of all DEGs) follows a similar pattern to what is observed across the entire genome for all the genes (49,5%). This indicates that the genes differently transcribed in F2-Mss(+) plants are not particularly enriched for TEs compared to other genes in the tomato genome. The transcription of TEs was also examined, using the RNAseq data, to determine whether TEs are deregulated in F2-Mss plants (adjusted p-value <0.05 and log2FoldChange <-1 or > 1). No TEs were found to be deregulated in F2-Mss(+) plants compared to the control driver lines and only one TE was downregulated in F2-Mss(-) plants. Employing identical bioinformatic procedures for comparison purposes, 7,783 TEs were found to be deregulated in *Slddm1* plants (Corem et al., 2018) with almost 92 % that were upregulated, which is in agreement with the function of DDM1 in promoting the maintenance of DNA methylation. Thus, when the bacterial methylase is expressed, only a very small fraction of TEs become deregulated in comparison to the total number of tomato TEs.

## DISCUSSION

Here, we expressed the bacterial CG-specific *M.SssI* methyltransferase under the control of the *CaMV 35S* promoter in tomato using the LhG4/pOP transactivation system, which separates the transformation and transgene expression steps. The plants expressing the methyltransferase are specifically hypermethylated in the CG context, in accessible chromatin regions, and thus mostly in genes. Conversely, heterochromatic regions are slightly hypomethylated in the CG context only. We also demonstrate that CG hyperDMRs produced by *M.SssI* can be inherited in the absence of bacterial methylase.

*M.SssI* fused to a ZF protein was introduced in *Arabidopsis* previously (Liu et al., 2021; Ghoshal et al., 2021). Even after multiple attempts, we were unsuccessful in directly introducing a *M.SssI* gene cloned in front of a double *CaMV 35S* promoter into *Arabidopsis* or tomato through transformation. Instead, the LhG4/pOP transactivation system (Moore et al., 1998) was successfully used in tomato. The variation with previous outcomes achieved in *Arabidopsis* may be attributed to the strength of the promoter used and therefore differences in *M.SssI* expression levels. Indeed, we employed a constitutive strong *CaMV 35S* promoter, while previous experiments were conducted with a *M.SssI*-ZF fusion driven by a *UBQ10* promoter (Liu et al., 2021), which is recognized for its ability to enable moderate expression in virtually all tissues of *Arabidopsis* (Grefen et al., 2010). An alternative explanation for the differences observed might be due to the experimental approaches. The *M.SssI*-ZF fusion protein exploited in *Arabidopsis* which was initially designed to target and bind to two neighboring repeats within the *FWA* promoter, demonstrated a broader binding capacity, affixing and functioning on numerous off-target sites. However, it is unclear whether the *M.SssI*-ZF fusion could bind without restriction to off-target sites or if those sites possess distinct features that are specifically recognized by the *M.SssI*-ZF fusion protein. The *M.SssI* used in this study was free of any fusion protein, therefore potentially preserving its capacity to bind a wider array of accessible target regions, potentially including novel targets, in comparison to the *M.SssI*-ZF fusion. Those targets may exhibit a heightened susceptibility to hypermethylation. In this study, we demonstrate that the *M.SssI* prokaryotic methyltransferase is active in tomato, a model crop, opening the door to targeted CG methylation as has already been demonstrated for *Arabidopsis* (Ghoshal et al., 2021) or mice embryos (Yamazaki et al., 2023). Moreover, our results suggest that a new type of epiRILs (Reinders et al., 2009; Johannes et al., 2009) can be generated by overmethylating DNA instead of stripping methylation, which could result in new epialleles and traits in crops. Plants expressing the bacterial methylase *M.SssI* show an overall modification of CG methylated sites. Methylation levels are more specifically increased within chromosome arms enriched for genes, and a small but significant decrease of CG methylation was detected in pericentromeric regions that are densely populated with repeats (Figure 3). In a recent study, we observed similar changes of DNA methylation homeostasis in the tomato *ddm1* mutants (Corem et al., 2018). DDM1 is a chromatin remodeler essential for maintaining DNA methylation and histone epigenetic marks, particularly in heterochromatic regions (Lyons and Zilberman, 2017; Lee et al., 2023b). In the *ddm1* mutant of tomato, the RdDM is partially redirected from euchromatin towards heterochromatin (Corem et al., 2018). Consequently, an imbalance in DNA methylation homeostasis occurred, marked by a reduction in both siRNAs and CHH methylation in chromosome arms and a parallel increase in heterochromatic regions. Thus, the RdDM pathway components appear to be diluted throughout the genome in *ddm1* tomato cells and certain elements of this pathway, such as enzymes or metabolites, may be limited in their availability. Other groups have obtained similar results with the *ddm1* mutants from rice (Tan et al., 2018). In this study, we expand upon this observation to show that the steady-state levels of CG methylation are also adjusted genome wide in tomato. Two possible hypotheses could explain the disturbance in CG methylation balance in plants expressing the bacterial CG-specific methylase. Firstly, the main endogenous enzyme responsible for maintaining CG methylation in plants, MET1, could be at limiting production to preserve the overall CG methylated sites including the one newly introduced by *M.SssI* along with the highly abundant CG methylated sites that are consistently present in heterochromatic regions. Secondly, the cell might not produce enough metabolites required by the DNA methyltransferases. The methylation of DNA requires S-adenosylmethionine, a universal methyl-group donor, as a cofactor, which is generated through the methionine cycle. *Arabidopsis* mutants impaired in the methionine cycle, like *mthfd1-1* (Groth et al., 2016), *methionine adenosyltransferase4* (Meng et al., 2018) or *methionine synthase1* (Yan et al., 2019) show decreased DNA and histone methylation, along with TE activation. It is therefore possible that S-adenosylmethionine is a limiting factor in *M.SssI* expressing plants, leading to the changes observed between CG methylation of euchromatin and heterochromatin when the bacterial methylase is active (Figure 3A and 3B).

Numerous studies have pointed out a modest correlation between changes in DNA methylation profiles and shifts of gene expression in plants (Goeldel and Johannes, 2023). The expression of the *M.SssI* bacterial methylase in tomato leads to a massive hypermethylation of genes in the CG context conducting to few changes in gene expression. Indeed, we found only 229 downregulated genes and 181 upregulated genes between plants expressing the methylase and the controls (Figure 6), suggesting that the changes in gene expression are not widespread across the genome. Still, an association between CG hypermethylation and transcriptional repression of genes was observed when genes were hypermethylated in the CG context near their TSS (Figure 7). Our findings revealed that genes exhibited a greater susceptibility to CG hypermethylation in the TSS region when they are expressed at lower levels in the wild type (Figure 7C). A recent study demonstrates that specific genes are particularly susceptible to alterations in CG gene body DNA methylation (Lee et al., 2023a). Loss of *DDM1* in *Arabidopsis* not only reduces DNA methylation, but also enhances resistance to a biotrophic pathogen when combined with mild chemical priming. The overall decrease in gene body methylation in the *ddm1* mutant additionally hyperactivates some stress-responsive genes leading to plant resistance (Lee et al., 2023a). Like many other genes, stress response genes are hypomethylated in a *ddm1* background but they become transcriptionally active only when the plants are attacked by a pathogen (Lee et al., 2023a). Therefore, modulating CG DNA methylation at specific genes weakly expressed might be a way to fine tune their regulation. The function of gene-body methylation, if any, remains enigmatic and our study extends to crops the observations of Liu *et al*. (Liu et al., 2021) by showing that global elevation of CG gene body methylation (Figure 3C) has minimal impact on the overall level of gene expression (Figure 6).

While we did observe a substantial quantity of CG hyperDMRs in F2-Mss(+) plants, those are somatic epimutations as DNAs analyzed were extracted from leaves. Transmission of the newly acquired methylation patterns to the next generations relies on the activity of *M.SssI* in the Shoot Apical Meristem (SAM), and more specifically in stem cells that serve as a functional germline. However, in tomato, the Arabidopsis *pFIL* promoter that we used to drive the expression of the *M.SssI* bacterial methyltransferase seems to be specific of leaf primordia, and significant expression within the SAM was not detected by using GUS or GFP reporters (Lifschitz et al., 2006). Nevertheless, we found that 20 to 32% of the CG hyperDMRs were transmitted between F2 plants expressing the methylase and their F3 progenies in which the transgene carrying the *M.SssI* gene is absent. Thus, CG methylation is likely transmitted to the SAM by an indirect mechanism that needs to be deciphered. Alternatively, the *pFIL* promoter might be active at very low rates in SAMs, possibly explaining why the number of DMRs transmitted to the next generation is lowered compared to *Arabidopsis M.SssI*-expressing plants where 50 to 90% of the DMRs are inherited (Liu et al., 2021). We also found that CG hyperDMRs newly appearing when the bacterial methylase is expressed are majorly not associated with CHG or CHH DMRs in both F2s and F3s. This is surprising considering that the RdDM pathway presumably triggers CHG and CHH methylation when methylation occurs (Zhou et al., 2018). Although they are few overlaps between CG hyperDMRs and DMRs found in other contexts, F2-Mss(+) CG hyperDMRs are slightly but significantly (t-test p-value < 0.001) hypermethylated in the CHG and CHH contexts (Figure 4D; Sup Figures 11 and 12). In the following F3-Mss(-) generation that inherited 20 to 30% of these CG hyperDMRs, the slight increase in both CHG and CHH methylation remains at similar levels, with no additional increase (Sup Figure 13). Therefore, we do not observe between F2 and F3 generations an enrichment of CG hyperDMRs in other forms of methylation. The effectiveness of RdDM might be hindered by a relatively low number of generations in our experiment. Alternatively, a recent study (Choi et al., 2021) demonstrated that the absence of CG methylation and histone H1 (*h1met1* mutants) led to a rise in methylation at CHH instead of the anticipated decrease. This finding suggests that CG methylation is not the crucial chromatin marker for the RdDM-dependent methylation observed at heterochromatic TEs in *h1* mutants. Instead, the authors suggest that CHG/CHH methylation serves as the main marker for attracting the RdDM machinery, with H3K9 methylation-dependent mechanisms playing a secondary role (Choi et al., 2021).

In conclusion, expressing the bacterial *M.SssI* methylase devoid of any fusion proteins via trans-activation has drastic consequences on the overall CG methylation homeostasis in tomato. CG DNA hypermethylation that is triggered in one generation through the expression of a foreign methyltransferase can be passed down to the subsequent generation. This opens possibilities for engineering precise DNA methylation in tomato plants. Activation of the *pOP*:*:M.SssI* responder line generated in this study by other driver lines with distinct cellular, tissue and organ specificities will facilitate the targeted modification of their epigenomes. This could further broaden the range of inherited epigenetic variations generated in this study. The resultant epigenetic variation could allow for the creation of new epiRIL populations. These populations have the potential to reveal previously unknown phenotypes that might not be identified solely by studying epigenetic variations in natural accessions.

## METHODS

### Plant material and growth conditions

The tomato (*S. Lycopersicum*) cv. M82 driver line *pFIL::LhG4* was described (Lifschitz et al., 2006). Germination and seedling growth took place in a growth chamber with a 16h light period and 8h dark period (photosynthetic photon flux density: 50 to 70 μmol m^-2^ s^-1^) at a constant temperature of 24°C. For crosses, closed flowers were emasculated by removal of the petals and stamens and hand-pollinated with the pollen of an appropriate homozygous driver line. Seeds were surface sterilized by treatment with 70% ethanol for 2 minutes followed by 3% sodium hypochlorite for 15 min. After rinsing three times with sterile distilled water, seeds were sown on MS culture medium with or without antibiotics. Germination and seedling growth were done in a growth chamber with a 16 h light/8 h dark period at a constant temperature of 24°C. Transgenic plants were moved and grown in 400 ml pots under greenhouse conditions with the temperature between 15-25°C in a peat mix with nutrients.

### Generation of tomato plants expressing *M.SssI*

Codon optimized bacterial methylase *M.SssI* with 2×35S promoter TMV omega, NLS and NOS terminator was synthesized (*Integrated DNA Technologies*, USA) and cloned into *pUC57* to create *pUC_M.SssI*. The potato *ST-LS1* IV intron (Eckes et al., 1986) was amplified (*Infusion*, Takara Bio) using *M.SssIN_IV* forward and *M.SssIC_IV* reverse primers (Sup Table 6) and cloned in the coding region of the methylase resulting in the *pUC_M.SssI_IV* plasmid carrying a disarmed plant adapted *M.SssI* (hereafter named *disM.SssI*). The methylase cassette was further subcloned in a binary vector *pART27* using the *Not*I restriction enzyme to yield *pART27_disM.SssI*. To clone under Op array, methylase was amplified using *M.SssI_HindIII* forward and reverse primers (Sup Table 6) cloned into *pGEMT-Easy* (*Promega*, USA) and sequenced for verification. The methylase cassette was further sub-cloned in a binary vector *pART27OP::P19HA* (Stav et al., 2010) vector digested with *HindIII* replacing *P19HA* to result in *pART27_OP::disM.SssI*.

The cotyledons of 14 days old tomato cv. M82 were transformed by co-cultivation with *Agrobacterium* strain GV3101 carrying the binary vector *pART27-OP::disMSssI* as described previously (Hendelman et al., 2013). Transgenic plants were selected on MS culture medium supplemented with Kanamycin (*Sigma*, USA). Presence of the transgene *pOP::disMSssI* was confirmed by PCR amplification on genomic DNA from plants that grew on selective media using *M.SssI-PCR* forward and *M.SssI-PCR* reverse primers (Sup Table 6). For crosses, pollen from the *pFIL:LhG4* diver line was used to hand pollinate the emasculated flower. F1 progenies were genotyped for the presence of *pOP::disMSssI* and *pFIL::LhG4* transgenes by PCR using *M.SssI-PCR* forward, *M.SssI-PCR* reverse primers and LhG4-forward, LhG4-reverse primers respectively (Sup Table 6).

### RNA analyses

Total RNA was extracted from 3 to 4 leaves of 45 days old plants, using Bio-Tri RNA reagent (*Bio-Lab*, Israel) according to the manufacturer’s instructions. DNase (*Ambion*, USA) treatment was performed on RNA samples to remove any residual genomic DNA. For qPCR analyses, 2 µg total RNA was used for first-strand cDNA synthesis using a Maxima first-strand cDNA synthesis kit (*Thermo Scientific*, USA) according to the manufacturer’s instructions. qRT-PCR was performed on the StepOne Plus real-time PCR system (*Applied Biosystems*, USA) using Fast SYBR Green Master Mix according to the manufacturer’s instructions. PCR products were analyzed using StepOne software version 2.2.2 (*Applied Biosystems*, USA). Expression levels were first normalized to a reference gene *TIP41* (SGN-U584254) (Lacerda et al., 2015) and relative expression levels were calculated using the 2^-ΔΔCt^ method.

For RNAseq, RNAs were extracted and treated with DNase as above. Total RNA was extracted from 3 to 4 leaves of 45 days old plants and the RNAs of these leaves were combined to sequence the transcriptome of each plant. Three plants (and therefore three biological replicates) were sequenced per genotype. Library preparation and sequencing were performed by *Macrogen* (Korea). On average, 38.4 million single-end 60 bp reads were sequenced per sample on HiSeq 2000 100 cycles run (Supplemental Table 5). The nf-core/RNAseq (version 3.11.0) pipeline (Ewels et al., 2020) was used for trimming (*TrimGalore*) and aligning (*STAR*) the reads to the SL2.5 version of the tomato genome assembly (The Tomato Genome Consortium, 2012) and to quantify reads matching transcripts (*Salmon*). The differential analysis was then performed with the obtained matrix of raw reads counts using the nf-core differential abundance pipeline (version 1.1.1) which is based on *DESeq2* (Love et al., 2014). RNAseq statistics are listed in Supplemental Table 5.

### Methylation analyses

To monitor the transient activity of *M.SssI* in tobacco leaves, *Agrobacterium tumefaciens* (strain GV 3101) with respective binary vectors were cultured overnight in LB medium containing the appropriate antibiotics. Cultures were pelleted and suspended in agroinfiltration buffer (10 mM MgCl_2_, 10 mM MES pH 5.6, and 150 µM acetosyringone) to a final O.D. at 600 of 1.0. Bacterial mixtures were then infiltrated into the young leaves of 3-week-old greenhouse-grown *N. benthamiana* plants. Methylation assays were performed after two days. Genomic DNA from leaves was extracted as described previously (Pavan Kumar et al., 2017). Global methylation (5-methyl cytosine (5mC)) was quantified Methylflash using 5- mC DNA ELISA Kit (*ZYMO Research*, USA) according to the manufacturer’s instructions.

To sequence the methylomes, DNA was extracted from 3 to 4 leaves of 45 days old plants with a genomic DNA extraction kit (*Macherey-Nagel*, England). The DNAs of these leaves were combined to sequence the methylome of each plant. Two to three plants (and therefore two to three biological replicates) were sequenced per genotype. Bisulfite treatments, library preparations, and whole-genome sequencings were performed at BGI (China) using HiSeq technology (Illumina), producing 150-bp paired-end reads. Data were trimmed with *Trim_Galore* (*Babraham Bioinformatics*). Reads were aligned to the SL2.5 tomato reference genome assembly (Tomato Genome Consortium, 2012) with *Bismark* version 0.22.3 (*Babraham Bioinformatics*) and standard options (*Bowtie2*; 1 mismatch allowed). Identical pairs were collapsed. To call Differently Methylated Regions (DMRs) between genotypes, we used the following R packages: *bsseq* version 1.30.0 (Hansen *et al*., 2012) and *DSS* version 2.42.0 (Wu *et al*., 2015). DMRs between the controls and other genotypes were identified considering the variation of each biological replicate. The following minimum thresholds were applied to define a DMR: 30% of difference for CG DMRs, 20% for CHG and 10% for CHH.

TEs were annotated with REPET (Flutre *et al*., 2011), and the repeat-rich, -intermediate and - poor regions were defined as described (Jouffroy *et al*., 2016), using the SL2.50 version of the genome.

Overlap between DMRs and accessible chromatin regions were determined genome-wide using available ATAC-seq (Assay for Transposase-Accessible Chromatin using sequencing) data of meristem-enriched tissue (Hendelman *et al*., 2021). Peak regions obtained by Hendelman *et al*. are accessible through GEO Series accession number GSE164297. Methylation and expression correlation were obtained with MethGet (Teng et al., 2020).

## SUPPLEMENTARY DATA

**Sup Figure 1.** Modified *M.SssI* methylation activity in *E. coli*.

**Sup Figure 2.** Transient expression of *M.SssI* in *Nicotiana benthamiana* leaves results in cytosine hypermethylation.

**Sup Figure 3.** Abnormal leaf phenotype observed in tomato plants overexpressing *disM.SssI*.

**Sup Figure 4.** Methylated cytosines (all contexts) in the plants analysed.

**Sup Figure 5.** Methylation along chromosomes, calculated from non-overlapping 200-kb bins.

**Sup Figure 6.** Correlation network diagram constructed using Spearman’s correlation coefficients to illustrate the relationships among the CG methylation bins depicted in the boxplots of Figure 3B.

**Sup Figure 7.** Patterns of methylation in genes in the four Mss-F2(+) plants expressing *disM.SssI* and the control drive lines.

**Sup Figure 8.** Patterns of methylation in Transposable Elements (TEs) in the four Mss-F2(+) plants expressing *disM.SssI* and the control drive lines.

**Sup Figure 9.** DMRs detected in F2 and F3 plants compared to the driver line control plants.

**Sup Figure 10**. Nature of the CG DMRs identified between the F2-Mss(+) plants and the driver line control plants.

**Sup Figure 11.** Patterns of methylation for CG hyperDMRs identified between F2-Mss(+) plant #1 and driver line controls.

**Sup Figure 12.** Patterns of methylation for CG hyperDMRs identified between F2-Mss(+) plant #3 and driver line controls.

**Sup Figure 13.** Patterns of methylation for CG hyperDMRs identified between F3-Mss(+) and driver line control plants.

**Sup Fig 14**. Principal Component Analysis (PCA) plots of RNAseq data.

**Sup Fig 15.** Methylation levels of genes that are up- or downregulated (> 1 ou <-1 log2FC) between the control and plants F2-Mss(+) expressing the methylase.

**Sup Fig 16.** Average methylation level profiling according to different expression groups around the TSS (+/- 200 bp) of F2-Mss(+) and control lines.

**Sup Table 1**. Whole-genome bisulfite sequencing statistics.

**Sup Table 2.** Number of DMRs identified in F2 and F3 plants compared to control driver lines.

**Sup Table 3.** RNAseq sequencing statistics.

**Sup Table 4:** List of upregulated genes between F2-Mss(+) plants and the control driver lines.

**Sup Table 5**: List of downregulated genes between F2-Mss(+) plants and the control driver lines.

**Sup Table 6:** Primers.

## ACKNOWLEDGEMENT

We thank Yoram Eyal and Isabelle Gy for helpful discussions and for critical reading of the manuscript.

We acknowledge funding from the Agence Nationale de la Recherche (Project ANR-21-CE20-0044, epiTOM) to NB, and Agricultural Research Organization manager funds to AS. The Institut Jean-Pierre Bourgin benefits from the support of the LabEx Saclay Plant Sciences-SPS (ANR-10-LABX-0040-SPS).

