## Supplemental Tables for "Genome wide inherited modifications of the tomato epigenome by trans-activated bacterial CG methyltransferase"

**Sup Table 1. Whole-genome bisulfite sequencing statistics**

| | Control driver lines | | | | F2-Mss(+) | | | | F2-Mss(-) | | F3-Mss(-) | |
|---|---|---|---|---|---|---|---|---|---|---|---|---|
| | **Plant#1** | **Plant#2** | **Plant#3** | **Plant#4** | **Plant#1** | **Plant#2** | **Plant#3** | **Plant#4** | **Plant#1** | **Plant#2** | **Plant#1** | **Plant#2** |
| Number of clean paired-end reads | 120,263,261 | 120,690,732 | 174,106,992 | 174,019,654 | 120,679,595 | 120,545,905 | 103,668,071 | 117,470,692 | 118,163,575 | 117,013,588 | 167,537,378 | 174,138,609 |
| Number of mapped paired-end reads | 96,773,350 | 98,764,278 | 138,591,199 | 137,439,583 | 79,997,757 | 81,876,303 | 74,533,989 | 90,921,225 | 86,014,361 | 82,464,462 | 132,537,678 | 137,723,956 |
| % of Cs methylated in the CG context | 77.4% | 78.0% | 72.3% | 74.4% | 78.1% | 79.4% | 77.7% | 77.3% | 78.7% | 75.3% | 80.2% | 78.4% |
| % of Cs methylated in the CHG context | 55.7% | 55.1% | 49.6% | 50.2% | 54.7% | 57.1% | 52.8% | 52.7% | 55.0% | 52.7% | 54.8% | 53.9% |
| % of Cs methylated in the CHH context | 9.1% | 9.4% | 7.4% | 6.8% | 11.5% | 12.4% | 11.6% | 11.3% | 11.9% | 10.5% | 5.9% | 6.8% |
| Coverage | 31.1 | 31.4 | 35.3 | 35.4 | 25.0 | 25.5 | 24.6 | 29.9 | 28.3 | 26.8 | 34.4 | 35.7 |
| Bisulfite conversion rates (%) | | | | | | | | | | | | |
| CG | 99.3% | 99.3% | 99.1% | 99.0% | 99.1% | 99.1% | 98.9% | 98.9% | 99.0% | 99.1% | 98.3% | 98.9% |
| CHG | 99.3% | 99.3% | 99.1% | 99.0% | 99.2% | 99.2% | 99.0% | 99.1% | 99.0% | 99.2% | 98.5% | 98.9% |
| CHH | 99.6% | 99.6% | 99.2% | 99.2% | 99.5% | 99.5% | 99.3% | 99.3% | 99.3% | 99.3% | 99.2% | 99.2% |

The number of clean paired-end reads obtained after trimming, the number of mapped paired-end reads, the coverage (after maping and deduplication), and the percentage of cytosines methylated in all contexts are indicated. Bisulfite conversion rates were determined by aligning the reads to the chloroplast sequences of *Solanum lycopersicum* (NCBI reference sequence NC_007898.3). The coverage was obtained using the *bam2nuc* script.

**Sup Table 2. Number of DMRs identified in F2 and F3 plants compared to control driver lines**

Total count and dimensions of DMRs for each genotype, and distribution of DMRs across regions with low TE density (RP), high TE density (RR), or intermediate TE presence (INT).

| | | | Count/size of DMRs (total) | | Distribution of DMRs | | |
|---|---|---|---|---|---|---|---|
| | | | number | DMR size (bp) | RP | INT | RR |
| F2-Mss(+) plant #1 | CG DMRs | hyper | 51795 | 281 | 31153 | 10184 | 11207 |
| | | hypo | 2650 | 176 | 1269 | 365 | 1040 |
| | CHG DMRs | hyper | 6076 | 235 | 1977 | 1517 | 2696 |
| | | hypo | 5809 | 206 | 952 | 858 | 4097 |
| | CHH DMRs | hyper | 40559 | 161 | 21399 | 10597 | 9296 |
| | | hypo | 1788 | 139 | 1012 | 418 | 390 |
| | | | | | RP | INT | RR |
| F2-Mss(+) plant #2 | CG DMRs | hyper | 48467 | 263 | 29135 | 9526 | 10513 |
| | | hypo | 2770 | 189 | 1260 | 379 | 1159 |
| | CHG DMRs | hyper | 7009 | 239 | 2244 | 1646 | 3239 |
| | | hypo | 7650 | 207 | 1585 | 1125 | 5069 |
| | CHH DMRs | hyper | 63586 | 177 | 33543 | 16138 | 15018 |
| | | hypo | 914 | 142 | 543 | 206 | 175 |
| | | | | | RP | INT | RR |
| F2-Mss(+) plant #3 | CG DMRs | hyper | 48254 | 277 | 28871 | 9483 | 10612 |
| | | hypo | 3745 | 195 | 2361 | 517 | 908 |
| | CHG DMRs | hyper | 5187 | 212 | 1799 | 1262 | 2219 |
| | | hypo | 8017 | 197 | 1784 | 1168 | 5187 |
| | CHH DMRs | hyper | 52247 | 165 | 26939 | 13201 | 13027 |
| | | hypo | 1109 | 139 | 652 | 241 | 238 |
| F2-Mss(+) plant #4 | CG DMRs | hyper | 54758 | 299 | 32574 | 10699 | 12272 |
| | | hypo | 3877 | 206 | 2092 | 531 | 1292 |
| | CHG DMRs | hyper | 5617 | 250 | 1712 | 1295 | 2702 |
| | | hypo | 11506 | 226 | 2241 | 1818 | 7629 |
| | CHH DMRs | hyper | 39870 | 165 | 21924 | 10076 | 8557 |
| | | hypo | 1214 | 166 | 562 | 292 | 381 |
| F2-Mss(-) plant #1 | CG DMRs | hyper | 12410 | 186 | 6867 | 2338 | 3382 |
| | | hypo | 745 | 176 | 434 | 107 | 208 |
| | CHG DMRs | hyper | 7289 | 226 | 2170 | 1538 | 3695 |
| | | hypo | 6313 | 222 | 1085 | 928 | 4404 |
| | CHH DMRs | hyper | 60537 | 169 | 32850 | 13954 | 14756 |
| | | hypo | 426 | 180 | 202 | 108 | 122 |
| F2-Mss(-) plant #2 | CG DMRs | hyper | 11177 | 180 | 6490 | 2176 | 2668 |
| | | hypo | 991 | 158 | 548 | 129 | 325 |
| | CHG DMRs | hyper | 6072 | 238 | 2160 | 1401 | 2607 |
| | | hypo | 8061 | 202 | 1259 | 1328 | 5636 |
| | CHH DMRs | hyper | 24425 | 145 | 13563 | 5603 | 5645 |
| | | hypo | 829 | 146 | 455 | 218 | 174 |

| | | | Count/size of DMRs (total) | | Distribution of DMRs | | |
|---|---|---|---|---|---|---|---|
| | | | number | DMR size (bp) | RP | INT | RR |
| F3-Mss(-) plant #1 | CG DMRs | hyper | 20354 | 186 | 13266 | 3638 | 3711 |
| | | hypo | 2760 | 147 | 1294 | 394 | 1112 |
| | CHG DMRs | hyper | 5273 | 204 | 2068 | 1252 | 2060 |
| | | hypo | 8253 | 203 | 1242 | 1189 | 5950 |
| | CHH DMRs | hyper | 370 | 138 | 218 | 74 | 83 |
| | | hypo | 25173 | 167 | 16035 | 5676 | 3889 |
| F3-Mss(-) plant #2 | CG DMRs | hyper | 13101 | 179 | 7690 | 2577 | 3056 |
| | | hypo | 4921 | 164 | 2392 | 775 | 1817 |
| | CHG DMRs | hyper | 5571 | 247 | 1593 | 1214 | 2860 |
| | | hypo | 9441 | 245 | 1748 | 1520 | 6319 |
| | CHH DMRs | hyper | 547 | 203 | 240 | 132 | 179 |
| | | hypo | 7957 | 146 | 4582 | 1913 | 1589 |

F2-Mss(+) plant #3 ---> F3-Mss(-) plant #1

F2-Mss(+) plant #4 ---> F3-Mss(-) plant #2

**Sup Table 3. RNAseq sequencing statistics**

| Sample | Total Bases | Read Count | GC (%) | AT (%) | Q20 (%) | Q30 (%) |
|---|---|---|---|---|---|---|
| F2Mss-plant3 | 4'088'370'718 | 40'478'918 | 43.52 | 56.48 | 98.15 | 94.34 |
| F2Mss-plant4 | 4'796'321'734 | 47'488'334 | 43.45 | 56.55 | 98.13 | 94.24 |
| F2Mss-plant5 | 5'040'808'192 | 49'908'992 | 43.49 | 56.51 | 98.12 | 94.23 |
| F2Mss+plant5 | 4'047'129'590 | 40'070'590 | 43.43 | 56.57 | 98.11 | 94.23 |
| F2Mss+plant6 | 3'898'406'686 | 38'598'086 | 43.35 | 56.65 | 98.26 | 94.53 |
| F2Mss+plant7 | 4'242'682'760 | 42'006'760 | 43.32 | 56.68 | 98.14 | 94.33 |
| controlplant5 | 3'665'476'244 | 36'291'844 | 43.45 | 56.55 | 98.05 | 94.05 |
| controlplant6 | 3'656'051'732 | 36'198'532 | 43.42 | 56.58 | 98.17 | 94.35 |
| controlplant7 | 3'747'230'290 | 37'101'290 | 43.13 | 56.87 | 98.19 | 94.40 |

**Sup Table 4**: List of upregulated genes between F2-Mss(+) plants and the control driver lines

| gene_id | baseMean | log2FoldChange | lfcSE | pvalue | padj |
|---|---|---|---|---|---|
| gene:Solyc12g100330.1 | 341.4361 | 1.002375 | 0.4818803 | 0.000308586 | 0.0189452 |
| gene:Solyc01g087720.2 | 487.0926 | 1.00739 | 0.26579115 | 3.32E-06 | 0.000694626 |
| gene:Solyc06g009680.2 | 208.2584 | 1.007704 | 0.36886821 | 6.78E-05 | 0.006863807 |
| gene:Solyc01g108350.2 | 15.84961 | 1.027854 | 1.79148372 | 0.000785007 | 0.03560864 |
| gene:Solyc04g071130.1 | 313.7964 | 1.03264 | 0.24052826 | 6.35E-07 | 0.000169584 |
| gene:Solyc07g056410.2 | 202.4333 | 1.042196 | 0.68665407 | 0.000959461 | 0.04053072 |
| gene:Solyc08g007670.1 | 151.8148 | 1.043352 | 0.45647182 | 0.000197617 | 0.01465398 |
| gene:Solyc12g044880.1 | 533.5657 | 1.045681 | 0.38979561 | 7.98E-05 | 0.007648742 |
| gene:Solyc06g036060.2 | 38.32235 | 1.046843 | 0.61356785 | 0.000667177 | 0.03196017 |
| gene:Solyc08g079860.1 | 466.6482 | 1.048283 | 0.78732436 | 0.001302062 | 0.04931359 |
| gene:Solyc09g065390.1 | 73.01785 | 1.052935 | 0.73889426 | 0.001114049 | 0.04473135 |
| gene:Solyc05g009650.2 | 406.5951 | 1.05953 | 0.69708383 | 0.000937266 | 0.03996381 |
| gene:Solyc04g076140.2 | 86.75811 | 1.062984 | 0.48389114 | 0.00024076 | 0.01638166 |
| gene:Solyc06g074500.1 | 98.88839 | 1.071848 | 0.35220561 | 3.42E-05 | 0.004099941 |
| gene:Solyc10g012210.1 | 40.53071 | 1.080888 | 0.58969792 | 0.000508013 | 0.026408 |
| gene:Solyc02g086580.2 | 52.65868 | 1.086803 | 0.53217283 | 0.000332335 | 0.01968054 |
| gene:Solyc02g077710.1 | 2300.296 | 1.109625 | 0.47755572 | 0.000184744 | 0.01409663 |
| gene:Solyc09g010690.2 | 964.6848 | 1.110837 | 0.58799417 | 0.000447068 | 0.02400077 |
| gene:Solyc07g055460.2 | 29.81812 | 1.119066 | 1.67540221 | 0.001010404 | 0.04216264 |
| gene:Solyc09g074230.2 | 529.2474 | 1.119497 | 0.65668913 | 0.000630105 | 0.0308407 |
| gene:Solyc02g091670.1 | 115.4466 | 1.136339 | 0.54692842 | 0.00030483 | 0.01876856 |
| gene:Solyc02g076800.1 | 164.7425 | 1.143546 | 0.37559961 | 3.75E-05 | 0.004442152 |
| gene:Solyc01g086660.2 | 84.40306 | 1.146791 | 0.8021526 | 0.000963707 | 0.04053072 |
| gene:Solyc03g007870.2 | 1194.59 | 1.148332 | 0.64634513 | 0.000538232 | 0.02764262 |
| gene:Solyc03g005570.2 | 293.5073 | 1.149689 | 0.88295574 | 0.001134876 | 0.04529992 |
| gene:Solyc07g055710.2 | 372.761 | 1.15314 | 0.76270598 | 0.000842896 | 0.03751764 |
| gene:Solyc01g095630.2 | 2118.204 | 1.155517 | 0.48717907 | 0.000167579 | 0.01301939 |
| gene:Solyc08g080650.1 | 5256.858 | 1.158938 | 1.06438095 | 0.001330622 | 0.04988775 |
| gene:Solyc06g051260.2 | 133.0599 | 1.159557 | 0.60357857 | 0.000407928 | 0.02276712 |
| gene:Solyc10g076510.1 | 179.8359 | 1.169531 | 0.92988801 | 0.001146458 | 0.04561281 |
| gene:Solyc12g005940.1 | 98.75981 | 1.170392 | 1.08980572 | 0.001304104 | 0.04931359 |
| gene:Solyc07g055080.2 | 245.5933 | 1.171982 | 0.46042724 | 0.000116979 | 0.01001291 |
| gene:Solyc05g052680.1 | 107.2752 | 1.176918 | 1.35628441 | 0.001260267 | 0.04861108 |
| gene:Solyc11g008440.1 | 212.1903 | 1.181095 | 0.32561025 | 8.81E-06 | 0.001471248 |
| gene:Solyc07g045100.1 | 341.8363 | 1.183055 | 0.61879414 | 0.000408135 | 0.02276712 |
| gene:Solyc12g009300.1 | 168.1554 | 1.190102 | 0.474398 | 0.000126605 | 0.01064927 |
| gene:Solyc11g008530.1 | 437.3991 | 1.199581 | 0.46371065 | 0.000107759 | 0.009320899 |
| gene:Solyc07g049550.2 | 256.5415 | 1.200334 | 0.94766625 | 0.001068924 | 0.04358315 |
| gene:Solyc12g010440.1 | 132.0414 | 1.20122 | 0.58046983 | 0.000300546 | 0.01873941 |
| gene:Solyc09g092760.2 | 109.6661 | 1.205579 | 0.77625022 | 0.00073432 | 0.03366682 |
| gene:Solyc03g114500.2 | 460.044 | 1.205601 | 0.82919938 | 0.000847666 | 0.03765154 |
| gene:Solyc06g066770.1 | 167.6796 | 1.207722 | 0.56881087 | 0.000270475 | 0.01745832 |
| gene:Solyc07g007170.2 | 463.0431 | 1.214159 | 0.5380335 | 0.000208826 | 0.01522717 |
| gene:Solyc11g066330.1 | 227.3807 | 1.214358 | 0.30589933 | 3.43E-06 | 0.000711972 |
| gene:Solyc04g007760.2 | 15.35227 | 1.220503 | 1.91289987 | 0.000585714 | 0.02951363 |

| | | | | | |
|---|---|---|---|---|---|
| gene:Solyc05g008540.2 | 20.38506 | 1.222135 | 0.56349274 | 0.000246239 | 0.01664843 |
| gene:Solyc06g066840.2 | 94.83042 | 1.223128 | 1.09543284 | 0.001136473 | 0.04529992 |
| gene:Solyc03g097610.2 | 646.6394 | 1.225107 | 0.5667608 | 0.000249264 | 0.01674694 |
| gene:Solyc08g007570.2 | 76.78254 | 1.22515 | 0.65239435 | 0.000418757 | 0.02305863 |
| gene:Solyc12g027540.1 | 11.45504 | 1.231962 | 0.9908425 | 0.001024353 | 0.04243358 |
| gene:Solyc01g105300.2 | 21.06099 | 1.23937 | 1.16653304 | 0.001118022 | 0.04473135 |
| gene:Solyc08g077210.2 | 855.6702 | 1.24551 | 0.32369256 | 5.22E-06 | 0.000979025 |
| gene:Solyc03g083480.2 | 623.7844 | 1.255706 | 0.9709417 | 0.000933751 | 0.03992278 |
| gene:Solyc05g015850.2 | 287.3131 | 1.260777 | 1.43585965 | 0.001034351 | 0.04265034 |
| gene:Solyc08g060920.2 | 293.3167 | 1.261932 | 0.55326075 | 0.00019536 | 0.01465398 |
| gene:Solyc03g117590.2 | 2543.916 | 1.264724 | 0.62833047 | 0.000318551 | 0.01924815 |
| gene:Solyc01g005500.2 | 13.18391 | 1.273688 | 1.51810137 | 0.000969128 | 0.04067864 |
| gene:Solyc12g055710.1 | 32.62982 | 1.282804 | 0.86243498 | 0.000713504 | 0.03328387 |
| gene:Solyc04g080520.1 | 95.69969 | 1.284858 | 0.38890759 | 2.29E-05 | 0.003151328 |
| gene:Solyc07g065320.2 | 2170.088 | 1.287802 | 0.43018932 | 4.63E-05 | 0.005180963 |
| gene:Solyc06g074360.2 | 12.70634 | 1.292278 | 1.07726303 | 0.000934303 | 0.03992278 |
| gene:Solyc06g071580.2 | 193.2214 | 1.295309 | 0.37245076 | 1.50E-05 | 0.002247309 |
| gene:Solyc02g081850.2 | 575.4082 | 1.300542 | 0.54441358 | 0.000156435 | 0.01247102 |
| gene:Solyc04g078540.2 | 636.6608 | 1.301683 | 0.22104942 | 9.54E-10 | 6.79E-07 |
| gene:Solyc04g064630.2 | 6.896174 | 1.301712 | 1.87464099 | 0.000621862 | 0.03061309 |
| gene:Solyc05g051780.2 | 151.8962 | 1.31247 | 0.25932937 | 6.65E-08 | 2.78E-05 |
| gene:Solyc01g079140.2 | 51.26915 | 1.318385 | 0.84560718 | 0.000612472 | 0.03043129 |
| gene:Solyc08g080190.2 | 120.1219 | 1.334821 | 0.72625536 | 0.000396872 | 0.02249116 |
| gene:Solyc09g011990.1 | 78.97583 | 1.336079 | 0.69382087 | 0.000344133 | 0.02025456 |
| gene:Solyc11g012930.1 | 825.6921 | 1.336799 | 0.76497945 | 0.000455228 | 0.02425425 |
| gene:Solyc09g007140.2 | 82.8581 | 1.364632 | 0.76616348 | 0.000416457 | 0.02304304 |
| gene:Solyc05g008210.2 | 108.2228 | 1.369643 | 0.96923437 | 0.000666795 | 0.03196017 |
| gene:Solyc01g103390.2 | 44.60026 | 1.372753 | 0.51656807 | 9.09E-05 | 0.008225197 |
| gene:Solyc05g009230.1 | 145.3765 | 1.38037 | 0.51321794 | 8.51E-05 | 0.007905729 |
| gene:Solyc12g009480.1 | 183.9734 | 1.394683 | 0.32917773 | 1.81E-06 | 0.000412214 |
| gene:Solyc09g082220.1 | 15.71283 | 1.397359 | 0.85107116 | 0.000483101 | 0.02542231 |
| gene:Solyc02g082920.2 | 19385.24 | 1.397654 | 1.10340217 | 0.000720151 | 0.0334619 |
| gene:Solyc10g008120.2 | 233.3956 | 1.403352 | 0.71254265 | 0.000296039 | 0.01860034 |
| gene:Solyc08g068610.2 | 211.3496 | 1.42303 | 1.13990461 | 0.000693915 | 0.03263811 |
| gene:Solyc09g073040.2 | 18.85446 | 1.428596 | 1.68036086 | 0.000682309 | 0.03243501 |
| gene:Solyc01g010060.2 | 155.1984 | 1.436939 | 0.97941012 | 0.000557016 | 0.02847043 |
| gene:Solyc02g067690.2 | 501.2118 | 1.451606 | 0.22245113 | 1.81E-11 | 2.76E-08 |
| gene:Solyc02g069690.1 | 160.5459 | 1.46089 | 0.48603161 | 4.39E-05 | 0.004963246 |
| gene:Solyc09g010360.2 | 364.2092 | 1.469498 | 0.72538704 | 0.000249231 | 0.01674694 |
| gene:Solyc04g077990.2 | 16.19825 | 1.500657 | 1.31125284 | 0.000630816 | 0.0308407 |
| gene:Solyc10g018120.1 | 85.7628 | 1.508703 | 0.94795874 | 0.000427108 | 0.02345802 |
| gene:Solyc09g018230.1 | 198.9432 | 1.515678 | 0.24223821 | 9.20E-11 | 1.03E-07 |
| gene:Solyc01g057270.2 | 20.15451 | 1.522087 | 0.58997779 | 9.37E-05 | 0.008344386 |
| gene:Solyc07g018200.1 | 56.24541 | 1.522373 | 1.42953376 | 0.000618312 | 0.03057926 |
| gene:Solyc05g056500.1 | 77.21236 | 1.528427 | 0.76274544 | 0.000237804 | 0.01628425 |
| gene:Solyc05g007770.2 | 585.809 | 1.530573 | 0.87012985 | 0.000334328 | 0.01973183 |
| gene:Solyc11g007200.1 | 1229.873 | 1.536664 | 0.38837435 | 4.31E-06 | 0.000845389 |
| gene:Solyc09g065120.2 | 5061.124 | 1.543965 | 0.52920703 | 4.93E-05 | 0.00537913 |

| | | | | | |
|---|---|---|---|---|---|
| gene:Solyc08g068850.2 | 26.91896 | 1.549709 | 0.75746135 | 0.000216757 | 0.015426 |
| gene:Solyc12g008680.1 | 18.61979 | 1.553526 | 0.70127388 | 0.00016626 | 0.012964 |
| gene:Solyc02g077770.2 | 119.9318 | 1.560893 | 0.43848574 | 1.22E-05 | 0.001893425 |
| gene:Solyc09g014990.2 | 778.9153 | 1.571364 | 0.70692211 | 0.000160509 | 0.01274822 |
| gene:Solyc10g007290.2 | 460.2584 | 1.574436 | 0.29311542 | 1.36E-08 | 7.45E-06 |
| gene:Solyc03g116480.1 | 106.3698 | 1.589383 | 0.85630177 | 0.000267596 | 0.01741634 |
| gene:Solyc08g014570.2 | 421.5918 | 1.600513 | 0.55484646 | 4.98E-05 | 0.005401335 |
| gene:Solyc10g086280.1 | 848.9063 | 1.623668 | 0.39971653 | 2.95E-06 | 0.000637694 |
| gene:Solyc01g112000.2 | 820.8798 | 1.62985 | 0.22551068 | 9.91E-14 | 3.53E-10 |
| gene:Solyc03g083590.2 | 50.06731 | 1.646239 | 0.36408 | 5.88E-07 | 0.000159669 |
| gene:Solyc12g088530.1 | 60.91296 | 1.650331 | 0.53855814 | 3.34E-05 | 0.00403282 |
| gene:Solyc09g009690.2 | 996.3906 | 1.656933 | 0.36285253 | 4.87E-07 | 0.000136802 |
| gene:Solyc11g012910.1 | 32.61335 | 1.660853 | 0.45117947 | 8.30E-06 | 0.00141802 |
| gene:Solyc10g076610.1 | 33.70978 | 1.672838 | 1.67916702 | 0.000462983 | 0.02454499 |
| gene:Solyc04g071800.2 | 839.0494 | 1.676861 | 0.25450766 | 7.74E-12 | 1.51E-08 |
| gene:Solyc05g056380.2 | 36.48751 | 1.677972 | 1.38332196 | 0.000441447 | 0.02381693 |
| gene:Solyc09g011610.2 | 14.99681 | 1.679391 | 1.17646975 | 0.000375809 | 0.02159213 |
| gene:Solyc11g012690.1 | 64.18936 | 1.6794 | 0.3282531 | 4.12E-08 | 1.83E-05 |
| gene:Solyc09g082230.1 | 219.4821 | 1.708112 | 0.58959238 | 4.30E-05 | 0.004907621 |
| gene:Solyc07g055690.1 | 36.04863 | 1.709204 | 1.00147283 | 0.000266047 | 0.01738258 |
| gene:Solyc01g110800.2 | 7.254863 | 1.730346 | 2.16068976 | 0.000285541 | 0.01815651 |
| gene:Solyc05g009610.1 | 410.0513 | 1.745885 | 0.62686612 | 5.01E-05 | 0.005402581 |
| gene:Solyc00g217960.1 | 64.89946 | 1.749212 | 0.49991927 | 1.17E-05 | 0.001832244 |
| gene:Solyc01g010230.2 | 321.1787 | 1.750238 | 0.17726569 | 8.43E-24 | 1.80E-19 |
| gene:Solyc08g079730.1 | 37.88215 | 1.771387 | 0.99977061 | 0.000222223 | 0.01561774 |
| gene:Solyc06g068500.2 | 2351.003 | 1.789686 | 0.68040782 | 6.15E-05 | 0.006379098 |
| gene:Solyc06g068940.2 | 9.63001 | 1.829029 | 1.4346218 | 0.000330555 | 0.01968054 |
| gene:Solyc12g098780.1 | 18.83397 | 1.837907 | 1.67548743 | 0.000350273 | 0.02050294 |
| gene:Solyc10g079420.1 | 1380.912 | 1.838235 | 0.38564575 | 1.50E-07 | 5.32E-05 |
| gene:Solyc01g100920.2 | 91.09602 | 1.839086 | 0.80787896 | 0.000100173 | 0.00884378 |
| gene:Solyc02g078160.2 | 296.5793 | 1.84651 | 0.64054871 | 3.64E-05 | 0.004338869 |
| gene:Solyc01g100200.2 | 1307.398 | 1.85263 | 0.401872 | 2.84E-07 | 8.78E-05 |
| gene:Solyc08g067760.2 | 13.22778 | 1.863546 | 1.0224399 | 0.000180123 | 0.01379326 |
| gene:Solyc11g072930.1 | 23.76639 | 1.895501 | 1.3514022 | 0.000269965 | 0.01745832 |
| gene:Solyc11g066250.1 | 496.1279 | 1.907025 | 0.31213889 | 8.88E-11 | 1.03E-07 |
| gene:Solyc09g011540.2 | 35.50011 | 1.908864 | 1.47204068 | 0.000287329 | 0.01819334 |
| gene:Solyc04g040120.1 | 110.7731 | 1.927784 | 1.74013236 | 0.000301821 | 0.01874537 |
| gene:Solyc03g093140.2 | 301.6011 | 1.981107 | 0.35661555 | 1.85E-09 | 1.27E-06 |
| gene:Solyc05g050130.2 | 6094.064 | 1.999872 | 0.67691532 | 2.50E-05 | 0.003377585 |
| gene:Solyc02g086960.2 | 7.881882 | 2.032759 | 2.02891123 | 0.000240437 | 0.01638166 |
| gene:Solyc05g046290.2 | 29.87052 | 2.041611 | 0.89453909 | 7.26E-05 | 0.007198936 |
| gene:Solyc03g111690.2 | 483.0454 | 2.043272 | 0.29996966 | 5.63E-13 | 1.50E-09 |
| gene:Solyc10g055810.1 | 9600.3 | 2.046617 | 0.54797265 | 3.83E-06 | 0.000779413 |
| gene:Solyc03g095770.2 | 1755.069 | 2.048809 | 0.4794499 | 6.63E-07 | 0.000174816 |
| gene:Solyc11g005350.1 | 413.5142 | 2.055888 | 0.63257788 | 1.25E-05 | 0.001913941 |
| gene:Solyc01g109680.2 | 66.46559 | 2.086773 | 0.37065419 | 8.52E-10 | 6.28E-07 |
| gene:Solyc01g106510.2 | 19.69522 | 2.090219 | 0.88885342 | 6.07E-05 | 0.006345879 |
| gene:Solyc10g007280.2 | 880.8283 | 2.091373 | 0.96813043 | 8.07E-05 | 0.007692915 |

| | | | | | |
|---|---|---|---|---|---|
| gene:Solyc02g094400.2 | 339.8682 | 2.104729 | 0.3892164 | 2.78E-09 | 1.80E-06 |
| gene:Solyc03g006490.2 | 1459.968 | 2.160998 | 0.85053606 | 4.02E-05 | 0.004668452 |
| gene:Solyc03g034390.1 | 16.40959 | 2.186989 | 1.75996272 | 0.000197009 | 0.01465398 |
| gene:Solyc02g068190.1 | 28.59458 | 2.307862 | 0.44968318 | 6.06E-09 | 3.50E-06 |
| gene:Solyc02g082740.1 | 287.9718 | 2.323964 | 0.70805345 | 7.11E-06 | 0.001254802 |
| gene:Solyc08g080620.1 | 2826.756 | 2.331035 | 2.31728725 | 0.00013626 | 0.01126646 |
| gene:Solyc12g094600.1 | 7.021757 | 2.341036 | 1.77088295 | 0.000153143 | 0.01227737 |
| gene:Solyc12g057090.1 | 84.08966 | 2.344528 | 0.52010821 | 1.11E-07 | 4.16E-05 |
| gene:Solyc03g005460.2 | 95.74972 | 2.362291 | 0.33974276 | 6.39E-14 | 3.41E-10 |
| gene:Solyc07g005380.2 | 45.23095 | 2.39995 | 1.69941298 | 0.000133254 | 0.01112102 |
| gene:Solyc07g042430.1 | 47.65533 | 2.412293 | 1.77849807 | 0.000136579 | 0.01126646 |
| gene:Solyc04g051490.2 | 83.53521 | 2.432591 | 0.33168643 | 3.06E-15 | 3.27E-11 |
| gene:Solyc09g098510.2 | 1360.506 | 2.438589 | 0.54958549 | 1.16E-07 | 4.26E-05 |
| gene:Solyc01g091490.2 | 77.30206 | 2.456278 | 0.58081252 | 2.65E-07 | 8.33E-05 |
| gene:Solyc03g036470.1 | 18.27031 | 2.456606 | 1.63143677 | 0.000114367 | 0.00985263 |
| gene:Solyc01g080900.2 | 58.95549 | 2.489741 | 0.38435811 | 1.06E-12 | 2.51E-09 |
| gene:Solyc00g026160.2 | 9901.052 | 2.50507 | 0.42845775 | 5.51E-11 | 7.85E-08 |
| gene:Solyc04g007000.1 | 596.096 | 2.527451 | 0.41909107 | 1.66E-11 | 2.72E-08 |
| gene:Solyc06g033860.1 | 92.66595 | 2.742751 | 1.2290513 | 3.14E-05 | 0.003853203 |
| gene:Solyc11g010670.1 | 34.02066 | 2.759153 | 0.58371991 | 1.31E-08 | 7.39E-06 |
| gene:Solyc03g025460.2 | 22.00318 | 2.775662 | 1.00597047 | 1.16E-05 | 0.001832244 |
| gene:Solyc12g056920.1 | 21.49372 | 2.957757 | 0.75022785 | 3.09E-07 | 9.42E-05 |
| gene:Solyc01g094380.2 | 36.74294 | 3.013659 | 1.13910634 | 1.14E-05 | 0.001817194 |
| gene:Solyc05g054010.2 | 51.64497 | 3.052154 | 0.90586896 | 1.94E-06 | 0.000435922 |
| gene:Solyc02g082910.2 | 51.19828 | 3.073165 | 1.47538262 | 2.88E-05 | 0.003726344 |
| gene:Solyc08g016270.1 | 44.90481 | 3.125969 | 0.79410232 | 2.60E-07 | 8.29E-05 |
| gene:Solyc01g010910.1 | 18.07336 | 3.160761 | 1.20894922 | 1.07E-05 | 0.001724265 |
| gene:Solyc03g006700.2 | 782.3427 | 3.306819 | 1.49751104 | 1.94E-05 | 0.002724976 |
| gene:Solyc01g099660.2 | 49.42795 | 3.311485 | 0.6770584 | 2.75E-09 | 1.80E-06 |
| gene:Solyc05g056400.2 | 61.54536 | 3.539661 | 1.61147338 | 1.63E-05 | 0.002376062 |
| gene:Solyc11g069880.1 | 28.69746 | 3.637082 | 1.90098585 | 2.17E-05 | 0.003007037 |
| gene:Solyc02g078150.2 | 384.3341 | 3.731784 | 0.94871705 | 1.99E-07 | 6.65E-05 |
| gene:Solyc02g086970.2 | 2861.048 | 3.970768 | 0.73529094 | 2.24E-10 | 1.99E-07 |
| gene:Solyc03g020030.2 | 76.41529 | 3.985491 | 1.4477253 | 4.70E-06 | 0.000888036 |
| gene:Solyc08g080590.2 | 256.654 | 4.000091 | 0.75717281 | 4.32E-10 | 3.69E-07 |
| gene:Solyc09g007940.2 | 55.22112 | 4.079544 | 1.59316999 | 6.20E-06 | 0.001112954 |
| gene:Solyc11g005480.1 | 117.7695 | 4.694104 | 1.37953011 | 8.73E-07 | 0.000219542 |
| gene:Solyc08g078180.1 | 80.80045 | 4.732774 | 1.53128823 | 1.63E-06 | 0.000373449 |
| gene:Solyc04g071070.2 | 849.151 | 4.935137 | 0.87840782 | 1.76E-10 | 1.71E-07 |
| gene:Solyc00g020840.2 | 199.1032 | 6.093411 | 1.68323007 | 1.95E-07 | 6.62E-05 |

**Sup Table 5**: List of downregulated genes between F2-Mss(+) plants and the control driver lines

| gene_id | baseMean | log2FoldChange | lfcSE | pvalue | padj |
|---|---|---|---|---|---|
| gene:Solyc07g042230.1 | 34.93491 | -6.170117 | 1.71736578 | 1.86E-07 | 6.52E-05 |
| gene:Solyc08g075210.1 | 194.0637 | -5.90794 | 0.85262144 | 9.54E-14 | 3.53E-10 |
| gene:Solyc01g095140.2 | 1353.009 | -5.44282 | 0.95959276 | 1.74E-10 | 1.71E-07 |
| gene:Solyc06g072670.2 | 32.37331 | -5.327921 | 1.85465578 | 1.39E-06 | 0.000330669 |
| gene:Solyc10g086690.1 | 51.98079 | -5.256407 | 1.44421978 | 3.95E-07 | 0.000112525 |
| gene:Solyc05g015800.2 | 57.24598 | -5.005243 | 1.3479308 | 3.72E-07 | 0.00010736 |
| gene:Solyc06g083900.2 | 181.871 | -4.72494 | 1.19665357 | 2.03E-07 | 6.66E-05 |
| gene:Solyc06g062370.2 | 1554.037 | -4.657158 | 0.76021972 | 7.76E-12 | 1.51E-08 |
| gene:Solyc11g071480.1 | 16.60675 | -4.554241 | 2.00658155 | 6.08E-06 | 0.001100915 |
| gene:Solyc08g083500.1 | 36.72113 | -4.39689 | 0.78365129 | 1.10E-10 | 1.17E-07 |
| gene:Solyc12g010970.1 | 49.29395 | -4.15693 | 1.19574361 | 8.43E-07 | 0.000214475 |
| gene:Solyc01g091170.2 | 2482.514 | -3.790216 | 0.72652859 | 5.25E-10 | 4.31E-07 |
| gene:Solyc04g012050.2 | 110.996 | -3.690598 | 1.35666137 | 5.95E-06 | 0.001087368 |
| gene:Solyc09g089540.2 | 37.84866 | -3.574106 | 2.3016802 | 2.93E-05 | 0.003745812 |
| gene:Solyc04g008330.1 | 53.10961 | -3.540656 | 1.20124036 | 4.03E-06 | 0.00080459 |
| gene:Solyc06g084190.2 | 9.140453 | -3.469278 | 1.96709973 | 2.97E-05 | 0.003776223 |
| gene:Solyc09g065620.2 | 34.17245 | -3.38018 | 1.00051994 | 1.45E-06 | 0.000341522 |
| gene:Solyc03g098790.1 | 449.8185 | -3.374082 | 1.27688846 | 8.55E-06 | 0.001438908 |
| gene:Solyc12g096780.1 | 57.57978 | -3.358162 | 0.98173301 | 1.30E-06 | 0.000312296 |
| gene:Solyc10g079980.1 | 21.25006 | -3.346901 | 1.44353916 | 1.57E-05 | 0.00229692 |
| gene:Solyc01g108800.2 | 76.63693 | -3.307857 | 0.51808027 | 4.07E-13 | 1.24E-09 |
| gene:Solyc02g085490.1 | 18.4365 | -3.220289 | 2.23106234 | 4.71E-05 | 0.005234732 |
| gene:Solyc01g088230.2 | 7.200877 | -3.186301 | 2.29229457 | 4.91E-05 | 0.005377892 |
| gene:Solyc08g065430.2 | 641.8166 | -3.150957 | 0.81843731 | 3.54E-07 | 0.000105126 |
| gene:Solyc03g007790.2 | 37.31878 | -3.096947 | 0.82054857 | 4.94E-07 | 0.000137095 |
| gene:Solyc08g066660.1 | 24.77879 | -3.012244 | 1.08056481 | 8.53E-06 | 0.001438908 |
| gene:Solyc10g080690.1 | 133.16 | -2.99801 | 1.39352231 | 2.77E-05 | 0.00361343 |
| gene:Solyc07g006570.2 | 875.1292 | -2.97051 | 0.945514 | 3.91E-06 | 0.000788914 |
| gene:Solyc02g069490.2 | 3232.822 | -2.939988 | 1.02004824 | 7.47E-06 | 0.001296961 |
| gene:Solyc05g055650.2 | 55.92656 | -2.927326 | 0.65223558 | 3.04E-08 | 1.51E-05 |
| gene:Solyc10g086500.1 | 571.528 | -2.902399 | 0.68792477 | 1.07E-07 | 4.16E-05 |
| gene:Solyc01g090560.2 | 174.8458 | -2.889932 | 0.6441675 | 3.26E-08 | 1.58E-05 |
| gene:Solyc08g007830.1 | 40.48235 | -2.888906 | 1.94630244 | 7.00E-05 | 0.007034428 |
| gene:Solyc11g005240.1 | 7.826506 | -2.879417 | 2.28392072 | 7.31E-05 | 0.007198936 |
| gene:Solyc03g111290.1 | 142.7603 | -2.776976 | 1.5867427 | 6.08E-05 | 0.006345879 |
| gene:Solyc09g007150.2 | 302.6652 | -2.709059 | 0.55063647 | 5.55E-09 | 3.29E-06 |
| gene:Solyc04g008370.1 | 87.31458 | -2.683646 | 0.82396017 | 4.17E-06 | 0.000824246 |
| gene:Solyc01g005870.1 | 1224.009 | -2.652314 | 0.90044144 | 9.11E-06 | 0.001497445 |
| gene:Solyc12g010960.1 | 42.58742 | -2.607082 | 1.17866945 | 3.76E-05 | 0.004442152 |
| gene:Solyc07g062930.2 | 76.65737 | -2.569997 | 0.47787313 | 6.87E-10 | 5.25E-07 |
| gene:Solyc04g054740.2 | 20392.61 | -2.565751 | 1.44866596 | 7.45E-05 | 0.007299861 |
| gene:Solyc03g013160.2 | 1987.563 | -2.546699 | 0.84390582 | 9.02E-06 | 0.001494141 |
| gene:Solyc09g083440.2 | 4933.715 | -2.535422 | 0.93685231 | 1.74E-05 | 0.002490019 |
| gene:Solyc09g008060.2 | 439.3945 | -2.491105 | 0.9211573 | 1.85E-05 | 0.002620155 |
| gene:Solyc04g051360.2 | 85.12711 | -2.404338 | 2.17631787 | 0.000139054 | 0.0114075 |

| | | | | | |
|---|---|---|---|---|---|
| gene:Solyc01g097520.2 | 5618.239 | -2.344192 | 0.66571516 | 3.72E-06 | 0.000765057 |
| gene:Solyc12g096770.1 | 782.1465 | -2.317902 | 1.22642728 | 8.58E-05 | 0.007940108 |
| gene:Solyc05g005960.2 | 1616.628 | -2.316231 | 0.51141111 | 1.10E-07 | 4.16E-05 |
| gene:Solyc08g083370.2 | 114.5749 | -2.281199 | 0.87588198 | 3.01E-05 | 0.003799451 |
| gene:Solyc10g079640.1 | 769.2232 | -2.26424 | 0.47507295 | 4.35E-08 | 1.90E-05 |
| gene:Solyc10g005360.2 | 80.78268 | -2.263403 | 0.55160693 | 7.10E-07 | 0.00018506 |
| gene:Solyc07g064600.2 | 2059.256 | -2.258538 | 1.04330933 | 6.30E-05 | 0.006467985 |
| gene:Solyc06g050520.1 | 5.943626 | -2.256092 | 1.84934732 | 0.000179883 | 0.01379326 |
| gene:Solyc01g090330.1 | 16.8826 | -2.252508 | 2.20775673 | 0.00016362 | 0.01292779 |
| gene:Solyc00g052940.2 | 121.0713 | -2.250968 | 1.02579118 | 6.09E-05 | 0.006345879 |
| gene:Solyc12g010980.1 | 1190.76 | -2.238213 | 1.41717143 | 0.000139357 | 0.0114075 |
| gene:Solyc03g078780.1 | 41.4898 | -2.237408 | 0.51973782 | 3.54E-07 | 0.000105126 |
| gene:Solyc08g005320.2 | 18.17883 | -2.23627 | 0.55275182 | 9.35E-07 | 0.000232206 |
| gene:Solyc12g010020.1 | 708.4305 | -2.227838 | 1.00328184 | 6.05E-05 | 0.006345879 |
| gene:Solyc02g087110.2 | 286.0134 | -2.227352 | 0.46084481 | 3.55E-08 | 1.65E-05 |
| gene:Solyc02g063000.2 | 10.02344 | -2.202973 | 1.73855027 | 0.000190461 | 0.0143788 |
| gene:Solyc06g071750.2 | 10.57135 | -2.198646 | 1.49417051 | 0.000164027 | 0.01292779 |
| gene:Solyc09g089510.2 | 2232.381 | -2.193412 | 1.11927183 | 9.23E-05 | 0.0082855 |
| gene:Solyc11g066670.1 | 657.9239 | -2.191161 | 1.24070059 | 0.000119195 | 0.01010558 |
| gene:Solyc02g084330.2 | 7.498982 | -2.148842 | 1.90677004 | 0.000212513 | 0.015339 |
| gene:Solyc07g055840.2 | 95.96163 | -2.140034 | 0.43308938 | 2.78E-08 | 1.42E-05 |
| gene:Solyc08g076970.2 | 11098.91 | -2.133693 | 0.28724661 | 4.70E-15 | 3.35E-11 |
| gene:Solyc01g096370.2 | 1089.548 | -2.12586 | 0.51953539 | 1.05E-06 | 0.000257654 |
| gene:Solyc02g085430.2 | 34.02353 | -2.094956 | 0.57596824 | 4.53E-06 | 0.000864531 |
| gene:Solyc06g074420.1 | 148.8634 | -2.088024 | 1.61175426 | 0.000220465 | 0.01554532 |
| gene:Solyc09g084490.2 | 1358.915 | -2.082077 | 1.75752827 | 0.000234313 | 0.01620099 |
| gene:Solyc03g098780.1 | 4359.588 | -2.077378 | 0.94686365 | 7.84E-05 | 0.007580676 |
| gene:Solyc02g094040.2 | 8.864983 | -2.07238 | 1.17264347 | 0.000140568 | 0.01141916 |
| gene:Solyc02g084770.2 | 35.08538 | -2.067884 | 0.4925938 | 8.33E-07 | 0.000214347 |
| gene:Solyc08g007210.2 | 67.40899 | -2.058926 | 0.50374974 | 1.25E-06 | 0.000304682 |
| gene:Solyc02g085730.2 | 3922.956 | -2.044465 | 0.63937308 | 1.42E-05 | 0.002139773 |
| gene:Solyc03g115370.2 | 563.4929 | -2.006413 | 0.52419681 | 3.20E-06 | 0.000675945 |
| gene:Solyc06g054240.2 | 21.80232 | -2.004694 | 0.7763801 | 4.79E-05 | 0.005277982 |
| gene:Solyc01g090340.2 | 483.8723 | -1.998535 | 1.7327207 | 0.000268195 | 0.01741634 |
| gene:Solyc11g022530.1 | 24.22839 | -1.933577 | 0.62346267 | 2.09E-05 | 0.002919409 |
| gene:Solyc06g007120.2 | 1209.011 | -1.924475 | 0.55707209 | 9.88E-06 | 0.001611027 |
| gene:Solyc07g054620.2 | 36.83931 | -1.894827 | 0.3875711 | 7.44E-08 | 3.00E-05 |
| gene:Solyc06g051940.2 | 408.8909 | -1.886243 | 1.09334927 | 0.000195673 | 0.01465398 |
| gene:Solyc02g086650.2 | 744.5886 | -1.878972 | 1.53635534 | 0.000314826 | 0.01916311 |
| gene:Solyc02g072440.2 | 21.91916 | -1.865463 | 1.46385398 | 0.000312067 | 0.01907616 |
| gene:Solyc01g105650.2 | 78.95627 | -1.862317 | 1.81150978 | 0.000332538 | 0.01968054 |
| gene:Solyc02g090340.2 | 263.6189 | -1.845182 | 0.69588633 | 5.47E-05 | 0.005868648 |
| gene:Solyc06g066540.1 | 151.8963 | -1.828853 | 1.19398894 | 0.000265318 | 0.01738258 |
| gene:Solyc02g091020.1 | 28.11219 | -1.818768 | 0.76801613 | 8.99E-05 | 0.008207358 |
| gene:Solyc03g093700.1 | 18.21884 | -1.814836 | 0.90538163 | 0.000153431 | 0.01227737 |
| gene:Solyc06g084100.2 | 5.590824 | -1.806488 | 1.80863244 | 0.000362318 | 0.02092139 |
| gene:Solyc03g098300.1 | 13.15242 | -1.770525 | 1.57470481 | 0.000390544 | 0.02219144 |
| gene:Solyc04g079730.1 | 690.4547 | -1.761608 | 1.32744964 | 0.000355604 | 0.02064533 |

| | | | | | |
|---|---|---|---|---|---|
| gene:Solyc03g078790.1 | 10.57402 | -1.758173 | 1.21170228 | 0.000322023 | 0.01932592 |
| gene:Solyc08g005350.2 | 195.813 | -1.752784 | 0.28104052 | 6.26E-11 | 8.36E-08 |
| gene:Solyc12g035710.1 | 78.77038 | -1.730006 | 0.42626068 | 2.63E-06 | 0.000585518 |
| gene:Solyc05g005950.2 | 3115.786 | -1.72578 | 0.56493671 | 3.11E-05 | 0.003853203 |
| gene:Solyc01g016370.1 | 40.52494 | -1.722699 | 1.28618068 | 0.000375955 | 0.02159213 |
| gene:Solyc07g053890.2 | 250.8708 | -1.720197 | 0.66516968 | 7.31E-05 | 0.007198936 |
| gene:Solyc06g063330.2 | 97.85574 | -1.70728 | 0.43789059 | 4.37E-06 | 0.000848095 |
| gene:Solyc12g098900.1 | 59.09244 | -1.704523 | 1.27057999 | 0.000387087 | 0.02211262 |
| gene:Solyc06g083040.2 | 5316.582 | -1.703828 | 0.77841519 | 0.000140135 | 0.01141916 |
| gene:Solyc02g068140.2 | 730.4193 | -1.690325 | 0.59161469 | 4.73E-05 | 0.005234732 |
| gene:Solyc03g062900.2 | 70.47614 | -1.689829 | 0.32986617 | 3.91E-08 | 1.78E-05 |
| gene:Solyc01g097390.2 | 77.00749 | -1.68963 | 0.45386362 | 7.22E-06 | 0.001263709 |
| gene:Solyc02g065240.2 | 86.06921 | -1.682544 | 0.29977824 | 3.04E-09 | 1.91E-06 |
| gene:Solyc02g071720.2 | 67.53885 | -1.680936 | 1.02834316 | 0.00030315 | 0.01876856 |
| gene:Solyc03g034460.2 | 33.51938 | -1.678023 | 0.54819611 | 3.26E-05 | 0.003956295 |
| gene:Solyc01g108880.2 | 92.91766 | -1.676886 | 0.41598086 | 3.07E-06 | 0.000655492 |
| gene:Solyc05g005920.2 | 897.7427 | -1.662424 | 0.31593771 | 2.06E-08 | 1.07E-05 |
| gene:Solyc12g088230.1 | 550.2587 | -1.650483 | 0.74876847 | 0.000148059 | 0.01198211 |
| gene:Solyc08g083230.1 | 61.43343 | -1.648519 | 0.55730911 | 4.12E-05 | 0.004753252 |
| gene:Solyc00g020540.1 | 10.65483 | -1.630986 | 1.85337004 | 0.000454456 | 0.02425425 |
| gene:Solyc02g070890.2 | 328.3922 | -1.626456 | 1.24983618 | 0.000457016 | 0.02428894 |
| gene:Solyc02g084350.2 | 63.76807 | -1.626209 | 0.83664511 | 0.000224136 | 0.0156715 |
| gene:Solyc01g087590.2 | 489.0451 | -1.623428 | 1.40594573 | 0.000499722 | 0.02604037 |
| gene:Solyc02g077420.2 | 658.1134 | -1.616436 | 0.7243521 | 0.00014898 | 0.01201117 |
| gene:Solyc06g068960.1 | 1095.099 | -1.611181 | 0.86382837 | 0.000255422 | 0.01696827 |
| gene:Solyc07g025170.2 | 75.0152 | -1.605872 | 0.51460797 | 3.12E-05 | 0.003853203 |
| gene:Solyc05g005930.1 | 433.7338 | -1.60049 | 0.30930044 | 3.54E-08 | 1.65E-05 |
| gene:Solyc03g013250.2 | 56.9294 | -1.590405 | 0.83863965 | 0.000252897 | 0.01693774 |
| gene:Solyc11g010170.1 | 1413.435 | -1.573862 | 0.59960599 | 8.21E-05 | 0.007699371 |
| gene:Solyc05g051460.2 | 595.9477 | -1.572181 | 0.56943271 | 6.45E-05 | 0.006588507 |
| gene:Solyc09g008640.1 | 362.3842 | -1.565259 | 0.32085032 | 1.47E-07 | 5.32E-05 |
| gene:Solyc10g008410.1 | 852.1116 | -1.557852 | 1.06022605 | 0.000447101 | 0.02400077 |
| gene:Solyc05g052470.2 | 660.661 | -1.552823 | 0.4343492 | 1.18E-05 | 0.001846871 |
| gene:Solyc03g116100.2 | 535.4516 | -1.552677 | 0.60153418 | 9.06E-05 | 0.008225197 |
| gene:Solyc05g005980.2 | 497.5385 | -1.551079 | 0.95636409 | 0.000383387 | 0.02195995 |
| gene:Solyc07g025160.2 | 238.104 | -1.543733 | 0.30642177 | 7.30E-08 | 3.00E-05 |
| gene:Solyc06g051060.2 | 27.05739 | -1.541856 | 0.51764356 | 4.39E-05 | 0.004963246 |
| gene:Solyc02g080120.1 | 185.6092 | -1.536173 | 1.08787907 | 0.000491068 | 0.02571484 |
| gene:Solyc12g057030.1 | 67.59374 | -1.51826 | 0.50341208 | 4.15E-05 | 0.004768273 |
| gene:Solyc08g078090.1 | 407.4595 | -1.515431 | 0.55411485 | 7.13E-05 | 0.007121381 |
| gene:Solyc03g116590.2 | 824.9216 | -1.508503 | 0.66849831 | 0.000164585 | 0.01292779 |
| gene:Solyc02g088790.2 | 346.7622 | -1.505312 | 0.2665533 | 3.40E-09 | 2.08E-06 |
| gene:Solyc07g005330.2 | 2404.137 | -1.504137 | 0.91922266 | 0.000408011 | 0.02276712 |
| gene:Solyc07g062700.2 | 14671.38 | -1.504018 | 0.99702543 | 0.000472632 | 0.0249328 |
| gene:Solyc01g110460.2 | 262.3901 | -1.501258 | 0.23850082 | 7.43E-11 | 9.33E-08 |
| gene:Solyc06g072430.1 | 76.1671 | -1.492915 | 0.90141975 | 0.000405665 | 0.02276712 |
| gene:Solyc12g042600.1 | 1065.715 | -1.491852 | 0.46070325 | 2.66E-05 | 0.003569254 |
| gene:Solyc10g081840.1 | 106.983 | -1.483154 | 0.89257588 | 0.000409453 | 0.02277233 |

| | | | | | |
|---|---|---|---|---|---|
| gene:Solyc01g096730.2 | 276.3153 | -1.482352 | 0.5550143 | 8.25E-05 | 0.007699371 |
| gene:Solyc06g068620.2 | 534.5748 | -1.46169 | 0.8133531 | 0.00035347 | 0.02063358 |
| gene:Solyc01g080910.2 | 645.2665 | -1.460105 | 0.63937881 | 0.000166177 | 0.012964 |
| gene:Solyc01g091320.2 | 961.4048 | -1.45412 | 1.275367 | 0.000692049 | 0.03263811 |
| gene:Solyc11g073080.1 | 35.05266 | -1.435503 | 0.6605298 | 0.000205077 | 0.01505663 |
| gene:Solyc03g006900.1 | 12.25034 | -1.435003 | 1.218401 | 0.000706639 | 0.03303577 |
| gene:Solyc12g036150.1 | 113.1738 | -1.430806 | 0.28844358 | 1.11E-07 | 4.16E-05 |
| gene:Solyc02g089010.1 | 39.41249 | -1.427912 | 0.80963708 | 0.000389841 | 0.02219144 |
| gene:Solyc07g007250.2 | 4204.205 | -1.409298 | 1.02414845 | 0.000642343 | 0.03126436 |
| gene:Solyc10g081570.1 | 1135.908 | -1.388034 | 1.02281316 | 0.000680699 | 0.03243501 |
| gene:Solyc11g006230.1 | 99.81088 | -1.38697 | 0.80030097 | 0.00043278 | 0.02358759 |
| gene:Solyc11g011570.1 | 103.9231 | -1.382722 | 0.39680746 | 1.54E-05 | 0.002280146 |
| gene:Solyc00g009130.2 | 47.70389 | -1.375958 | 0.36025447 | 6.41E-06 | 0.001140472 |
| gene:Solyc01g081160.2 | 11.62794 | -1.374455 | 1.00636273 | 0.000692833 | 0.03263811 |
| gene:Solyc01g094010.2 | 839.5902 | -1.370081 | 0.70345595 | 0.000319172 | 0.01924815 |
| gene:Solyc01g107800.2 | 125.3672 | -1.350678 | 0.75363229 | 0.000417396 | 0.02304304 |
| gene:Solyc11g005430.1 | 31.32725 | -1.347342 | 1.16329178 | 0.000852851 | 0.03780322 |
| gene:Solyc02g071380.2 | 109.5405 | -1.344627 | 0.65547318 | 0.00027665 | 0.01774965 |
| gene:Solyc06g035940.2 | 265.7163 | -1.339991 | 1.47082968 | 0.000879283 | 0.03873376 |
| gene:Solyc12g036140.1 | 226.5248 | -1.33899 | 0.24910485 | 1.52E-08 | 8.11E-06 |
| gene:Solyc01g099290.2 | 77.59252 | -1.337573 | 0.35952548 | 8.23E-06 | 0.00141802 |
| gene:Solyc06g065910.2 | 8.501644 | -1.336135 | 1.301642 | 0.000907999 | 0.03954841 |
| gene:Solyc12g014450.1 | 279.6693 | -1.332377 | 0.293451 | 5.90E-07 | 0.000159669 |
| gene:Solyc01g105890.2 | 54.38722 | -1.31666 | 0.8691858 | 0.000650789 | 0.03152857 |
| gene:Solyc07g048060.1 | 60.2612 | -1.314048 | 0.91803983 | 0.000723432 | 0.0334619 |
| gene:Solyc08g075790.2 | 205.9035 | -1.309971 | 0.63967653 | 0.000287824 | 0.01819334 |
| gene:Solyc10g086250.1 | 106.8644 | -1.301925 | 1.30302763 | 0.000979258 | 0.04102323 |
| gene:Solyc08g074630.1 | 259.6487 | -1.297745 | 1.0922348 | 0.000930424 | 0.03992278 |
| gene:Solyc12g049350.1 | 21.08583 | -1.294812 | 1.59076208 | 0.000891832 | 0.03912522 |
| gene:Solyc06g062960.1 | 237.9816 | -1.294118 | 0.58473093 | 0.000217329 | 0.015426 |
| gene:Solyc09g072710.2 | 308.4399 | -1.290344 | 0.3659893 | 1.33E-05 | 0.002025766 |
| gene:Solyc02g087800.2 | 55.67929 | -1.289602 | 0.47307222 | 8.12E-05 | 0.007699371 |
| gene:Solyc11g020820.1 | 148.3351 | -1.288386 | 0.3344741 | 5.46E-06 | 0.001005243 |
| gene:Solyc03g123540.2 | 14.8948 | -1.288348 | 1.03791155 | 0.000914332 | 0.03962415 |
| gene:Solyc09g007260.2 | 462.3665 | -1.28217 | 0.86791828 | 0.000723585 | 0.0334619 |
| gene:Solyc07g061800.2 | 423.815 | -1.27886 | 1.54703606 | 0.000943433 | 0.04007245 |
| gene:Solyc08g069140.2 | 342.053 | -1.273874 | 0.39557032 | 2.77E-05 | 0.00361343 |
| gene:Solyc05g024160.2 | 209.5671 | -1.26104 | 0.41048121 | 3.86E-05 | 0.004519288 |
| gene:Solyc09g015000.2 | 33.35463 | -1.254792 | 0.56429135 | 0.000218339 | 0.01544637 |
| gene:Solyc03g007240.2 | 1336.116 | -1.240466 | 0.4650315 | 9.16E-05 | 0.008255814 |
| gene:Solyc01g091360.2 | 416.2305 | -1.240458 | 0.56123715 | 0.000225614 | 0.01570112 |
| gene:Solyc12g006460.1 | 1748.641 | -1.233822 | 0.80342738 | 0.000722635 | 0.0334619 |
| gene:Solyc01g009570.2 | 161.1153 | -1.22713 | 0.77503494 | 0.00068013 | 0.03243501 |
| gene:Solyc10g052490.1 | 320.5179 | -1.226047 | 0.82654497 | 0.000787604 | 0.03565076 |
| gene:Solyc05g041420.2 | 193.8574 | -1.224483 | 0.20575771 | 6.11E-10 | 4.84E-07 |
| gene:Solyc01g098240.1 | 11.57727 | -1.22358 | 0.9806521 | 0.001038061 | 0.04265034 |
| gene:Solyc04g081290.2 | 846.1981 | -1.216566 | 0.65136181 | 0.000429766 | 0.02348324 |
| gene:Solyc10g008840.2 | 113.4682 | -1.209975 | 0.59507888 | 0.000319827 | 0.01924815 |

| | | | | | |
|---|---|---|---|---|---|
| gene:Solyc03g007340.2 | 11.83654 | -1.19983 | 1.42756952 | 0.001164485 | 0.04615809 |
| gene:Solyc08g066450.1 | 52.7802 | -1.188142 | 0.5342064 | 0.000224455 | 0.0156715 |
| gene:Solyc00g009120.2 | 50.54036 | -1.183265 | 0.43650762 | 8.25E-05 | 0.007699371 |
| gene:Solyc12g076320.1 | 36.93039 | -1.181 | 0.40899266 | 5.58E-05 | 0.005964039 |
| gene:Solyc01g058260.2 | 1802.874 | -1.179699 | 0.37018295 | 2.77E-05 | 0.00361343 |
| gene:Solyc06g082030.2 | 26.09326 | -1.175469 | 0.81066897 | 0.00089456 | 0.0391645 |
| gene:Solyc07g042540.2 | 10.52374 | -1.169047 | 1.20581097 | 0.001330963 | 0.04988775 |
| gene:Solyc10g050450.1 | 11.16551 | -1.163342 | 0.89741938 | 0.001113468 | 0.04473135 |
| gene:Solyc08g013820.2 | 57.15493 | -1.153083 | 0.46369227 | 0.000131863 | 0.01104807 |
| gene:Solyc03g098760.1 | 19.8772 | -1.152156 | 1.85199441 | 0.00068468 | 0.03243501 |
| gene:Solyc03g122270.2 | 1307.41 | -1.144505 | 0.18725758 | 1.98E-10 | 1.84E-07 |
| gene:Solyc07g054630.2 | 48.3777 | -1.141225 | 0.57100758 | 0.000355382 | 0.02064533 |
| gene:Solyc03g118540.2 | 49.10217 | -1.135135 | 0.58686744 | 0.000403968 | 0.02276712 |
| gene:Solyc03g121100.2 | 13.76043 | -1.134242 | 0.9055306 | 0.001235717 | 0.04835366 |
| gene:Solyc02g064620.2 | 305.8175 | -1.127755 | 0.43141339 | 9.94E-05 | 0.008808819 |
| gene:Solyc02g091030.2 | 404.723 | -1.126931 | 0.35899937 | 2.92E-05 | 0.003745812 |
| gene:Solyc06g005910.2 | 141.1927 | -1.120013 | 0.66350171 | 0.000649319 | 0.03152857 |
| gene:Solyc06g076670.2 | 710.1365 | -1.115025 | 0.64822387 | 0.00061502 | 0.03048699 |
| gene:Solyc06g066650.2 | 218.9014 | -1.112457 | 0.39247488 | 5.97E-05 | 0.006343678 |
| gene:Solyc01g080640.2 | 4672.287 | -1.104818 | 0.49152336 | 0.000216607 | 0.015426 |
| gene:Solyc06g068700.2 | 21.53383 | -1.099495 | 0.62482956 | 0.000578381 | 0.02937373 |
| gene:Solyc01g098320.2 | 104.8029 | -1.091427 | 0.39203526 | 6.58E-05 | 0.006694976 |
| gene:Solyc05g010540.2 | 126.0298 | -1.087954 | 0.2769539 | 2.83E-06 | 0.000622367 |
| gene:Solyc03g097170.2 | 105.0364 | -1.084961 | 0.68752644 | 0.000820991 | 0.0369273 |
| gene:Solyc03g083620.1 | 632.2912 | -1.077305 | 0.79111526 | 0.001188146 | 0.0469219 |
| gene:Solyc07g063910.2 | 404.4276 | -1.07573 | 0.80796968 | 0.001245199 | 0.04854686 |
| gene:Solyc02g091140.2 | 849.4633 | -1.071772 | 0.34787104 | 3.10E-05 | 0.003853203 |
| gene:Solyc02g085870.2 | 4184.265 | -1.07062 | 0.32248372 | 1.65E-05 | 0.002377221 |
| gene:Solyc12g099000.1 | 2923.397 | -1.064723 | 0.64450878 | 0.000730489 | 0.03363555 |
| gene:Solyc02g088770.2 | 54.51068 | -1.060455 | 0.5004741 | 0.000284224 | 0.01812672 |
| gene:Solyc00g006820.2 | 1078.232 | -1.059627 | 0.70290344 | 0.000958153 | 0.04053072 |
| gene:Solyc09g031790.2 | 120.1469 | -1.058097 | 0.48341027 | 0.000244783 | 0.01660248 |
| gene:Solyc01g091850.2 | 33.7035 | -1.0569 | 0.79030536 | 0.001272961 | 0.04873982 |
| gene:Solyc02g070530.2 | 256.275 | -1.037628 | 0.4540419 | 0.00019742 | 0.01465398 |
| gene:Solyc11g066680.1 | 316.7362 | -1.023456 | 0.73833495 | 0.001238136 | 0.04835975 |
| gene:Solyc04g074270.2 | 2170.921 | -1.02243 | 0.50930862 | 0.000361436 | 0.02092139 |
| gene:Solyc01g080700.2 | 223.9018 | -1.020267 | 0.4742587 | 0.000264354 | 0.0173782 |
| gene:Solyc02g068030.1 | 755.733 | -1.019998 | 0.7490701 | 0.001296002 | 0.04931359 |
| gene:Solyc12g036120.1 | 80.67339 | -1.004936 | 0.3641058 | 6.30E-05 | 0.006467985 |
| gene:Solyc12g027800.1 | 84.26658 | -1.004643 | 0.33313779 | 3.26E-05 | 0.003956295 |

**Sup Table 6: Primers**

| Primer name | Sequence (5'-3') | Purpose |
|---|---|---|
| MSssI_PCR_For | AAGTTGAGAACAAGACTAAGAA | Detection of methylase gene in transgenic plants |
| MSssI_PCR_Rev | GTTCTTGAGAGTTCTCTTATAA | |
| MSssIN-IV-R | AGGTAGAAGCAGAAACTTACCTGCGAATGCTTCGAACAC | Cloning of the potato *IV2* intron |
| MSssIN-IV-F | GTGTTCGAAGCATTCGCAGGTAAGTTTCTGCTTCTACCT | |
| MSssIC-IV-R | CTTTTCTTTGTGCACCTATACCTGCACATCAACAAATTTTG | |
| MSssIC-IV-F | CAAAATTTGTTGATGTGCAGGTATAGGTGCACAAAGAAAAG | |
| MSssI_HindIII For | CGCAAGCTTATGTCTAAAGTTGAGAACAAG | Cloning of methylase |
| MSssI_HindIII Rev | CGCAAGCTTTCACACCTTCCTCTTCTTC | |
| Lhg4_453_FWD | TGACCAGACACCCATCAACAGTA | Detection of *Lhg4* transgene |
| Lhg4_1004_rev | GGCGAATTCCGATCCACTAG | |
| qRT-TIP41-Fwd | ATGGAGTTTTTGAGTCTTCTGC | qRT-PCR |
| qRT-TIP41-Rev | GCTGCGTTTCTGGCTTAGG | |
| qRT-MSssI-Fwd | AGCATTCGCAGGTATAGGTGC | qRT-PCR |
| qRT-MSssI-Rev | CTTCCAATATCCGTTAGACACAG | |
