## Supplemental Figures for "Genome wide inherited modifications of the tomato epigenome by trans-activated bacterial CG methyltransferase"

Sup Figure 1

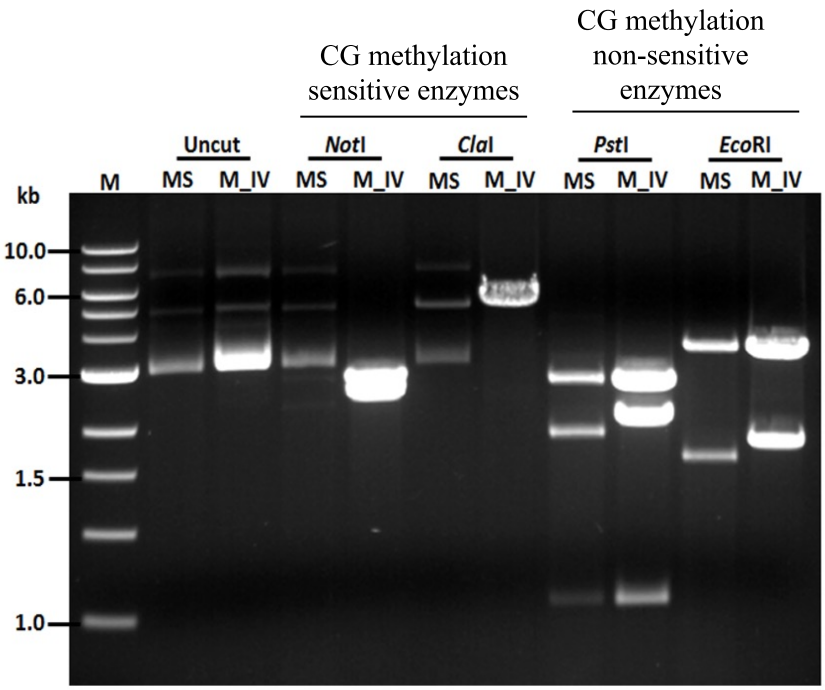


**Sup Figure 1. Modified *M.SssI* methylation activity in *E. coli***

Loss of restriction sites sensitive to methylation was tested on plasmids in *E. coli* expressing *M.SssI*. The DNA was digtested by CG methylation sensitive (*ClaI*, *NotI*) and non-sensitive (*PstI*, *EcoRI*) enzymes. MS: *pUC::M.SssI*, a plasmid expressing the methylase with a *35S* promoter and M_IV: *pUC::disM.SssI* plasmid expressing the disarmed methylase with the potato *IV2* intron (inactive in *E. coli*).

MS is not cut by *NotI* and *ClaI* (methylation sensitive enzymes) but is cut by *PstI* and *EcoRI* (methylation non-sensitive enzymes).

*M*, Marker.

Sup Figure 2

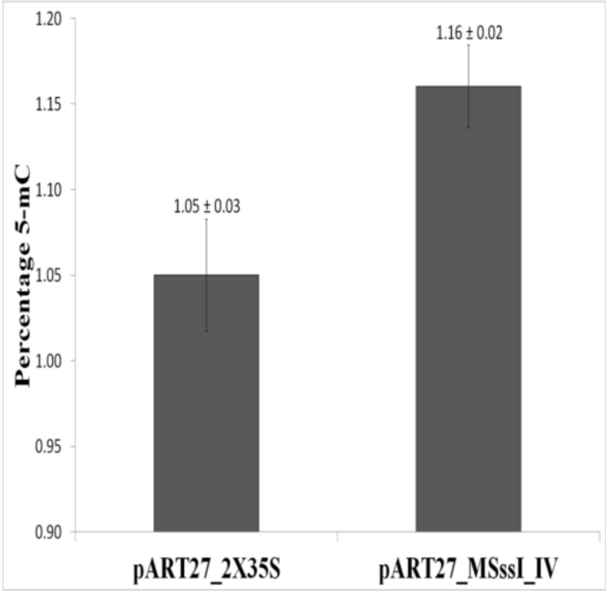


**Sup Figure 2**. **Transient expression of *M.SssI* in *Nicotiana benthamiana* leaves results in cytosine methylation**

Quantification of global 5-methyl cytosine in the genomic DNA of *Nicotiana benthamiana* leaves two days post infiltration with *pART27_disM.SssI* and *pART27_2X35S* used as control. Error bars indicate ±SD over two biological replicates.

Sup Figure 3

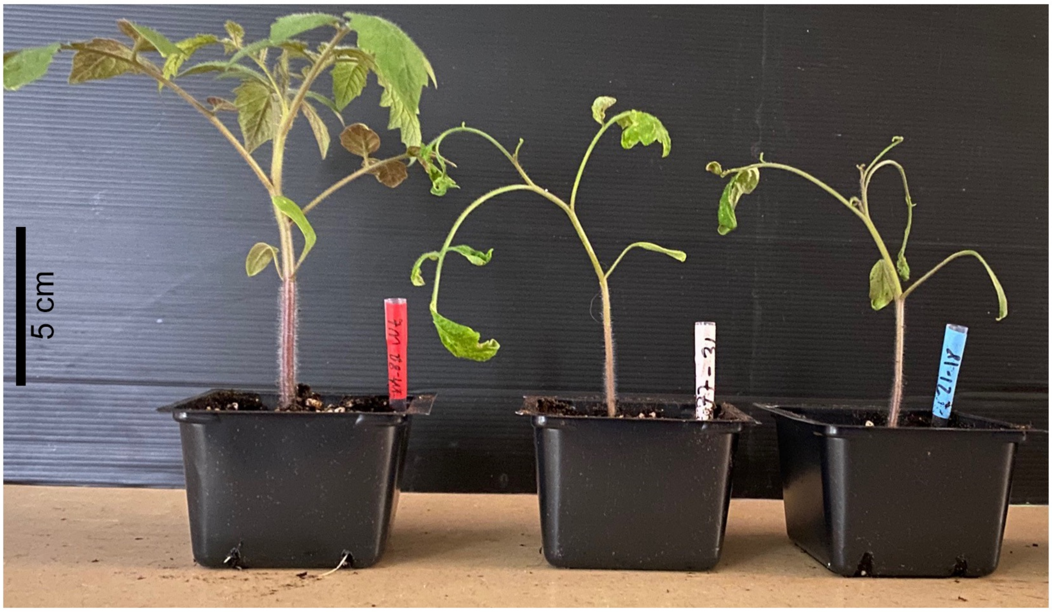


M82 plant

*pFIL::LhG4* >> p*OP::disM.SssI*
F3 plants

**Sup Figure 3. Abnormal leaf phenotype observed in tomato plants overexpressing *disM.SssI*.**

F3 plants exhibiting the abnormal leaf phenotype and M82 used as control were pictured 30 days after germination.

The phenotype consistently manifests in plants that overexpress M.SssI, although full penetrance is not achieved, as evidenced by the fact that not all plants carrying the activation system display the phenotype. Indeed, investigations into F3 progenies of an F2 plant double heterozygous for the two constructs and showing the leaf phenotype revealed that 10 out of 54 plants exhibited the phenotype. Conversely, in F3 progenies of an F2 plant lacking the M.SssI activation system, the phenotype was completely absent, with none of the 50 progeny plants showing it.

Sup Figure 4

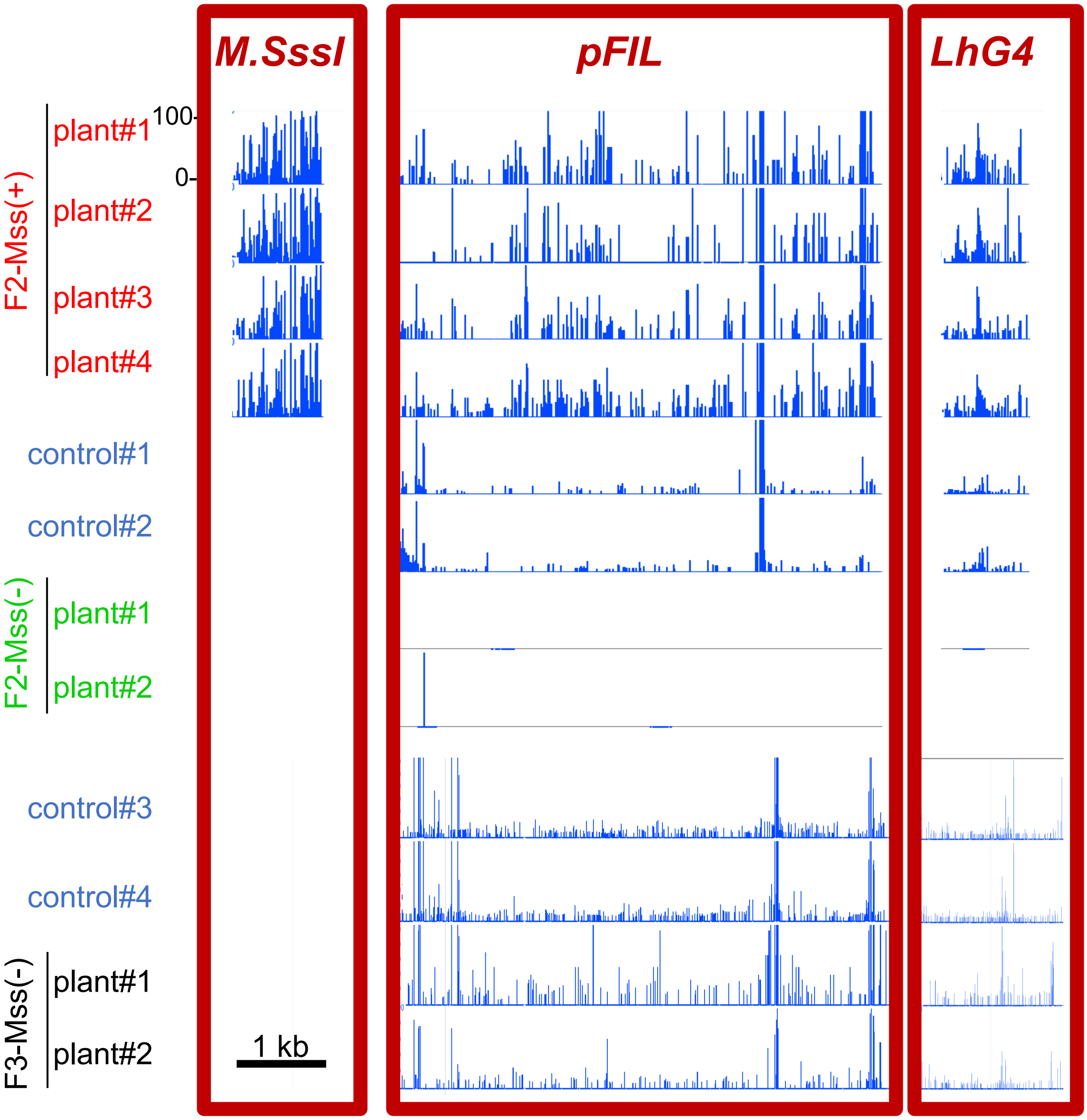


**Sup Figure 4**. Methylated cytosines (CG, CHG and CHH methylation contexts are plotted together) in the plants analysed.
Reads were aligned to the three transgene components: the *M.SssI* gene, the *pFIL* promotor and the *LhG4* gene. The methylation levels are given in % (scale: 0 to 100%).

### Sup Figure 5

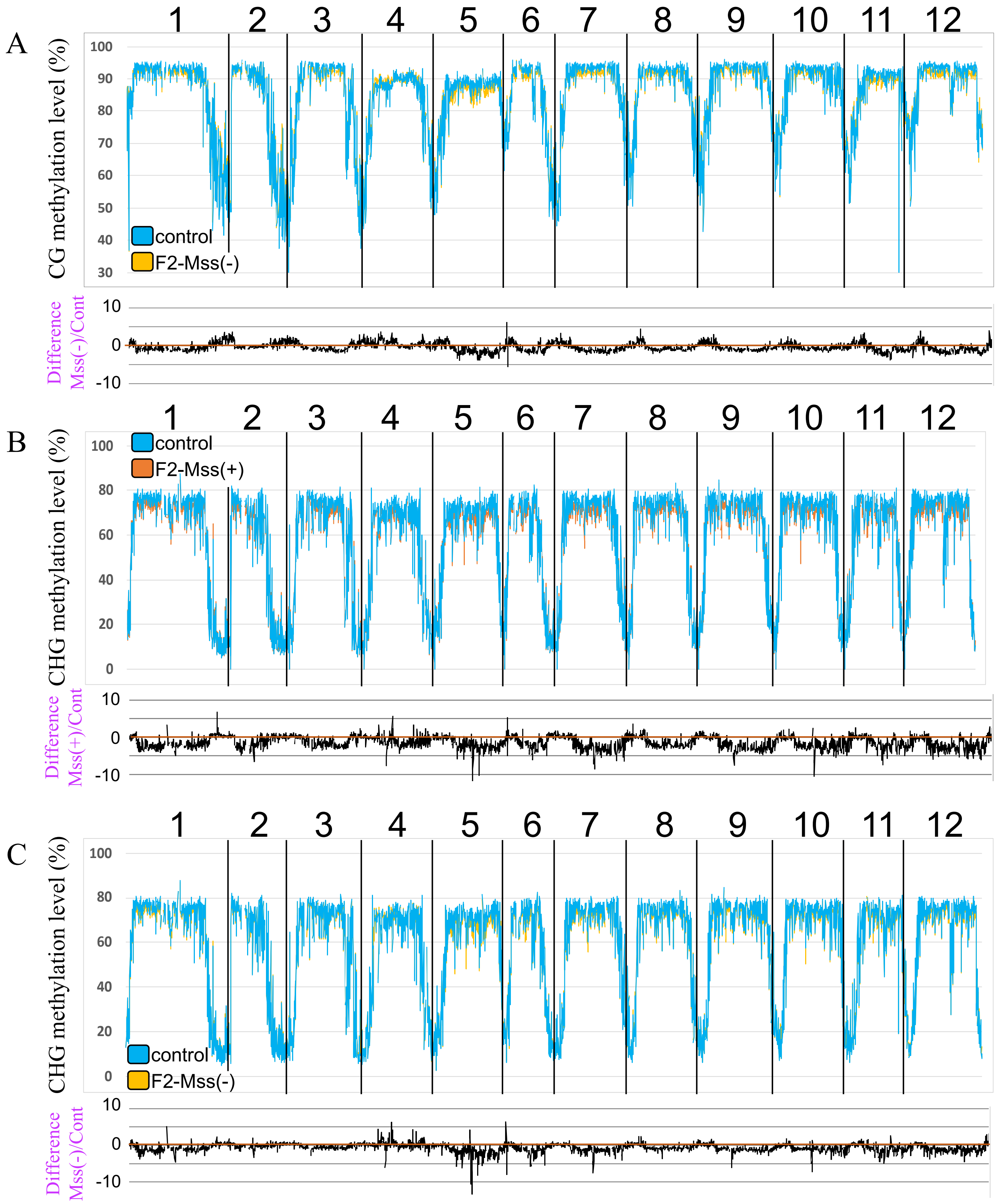
A
1 2 3 4 5 6 7 8 9 10 11 12
CG methylation level (%)
100 90 80 70 60 50 40 30
control
F2-Mss(-)
Difference Mss(-)/Cont
10 0 -10
B
1 2 3 4 5 6 7 8 9 10 11 12
CHG methylation level (%)
100 80 60 40 20 0
control
F2-Mss(+)
Difference Mss(+)/Cont
10 0 -10
C
1 2 3 4 5 6 7 8 9 10 11 12
CHG methylation level (%)
100 80 60 40 20 0
control
F2-Mss(-)
Difference Mss(-)/Cont
10 0 -10

Sup Figure 5

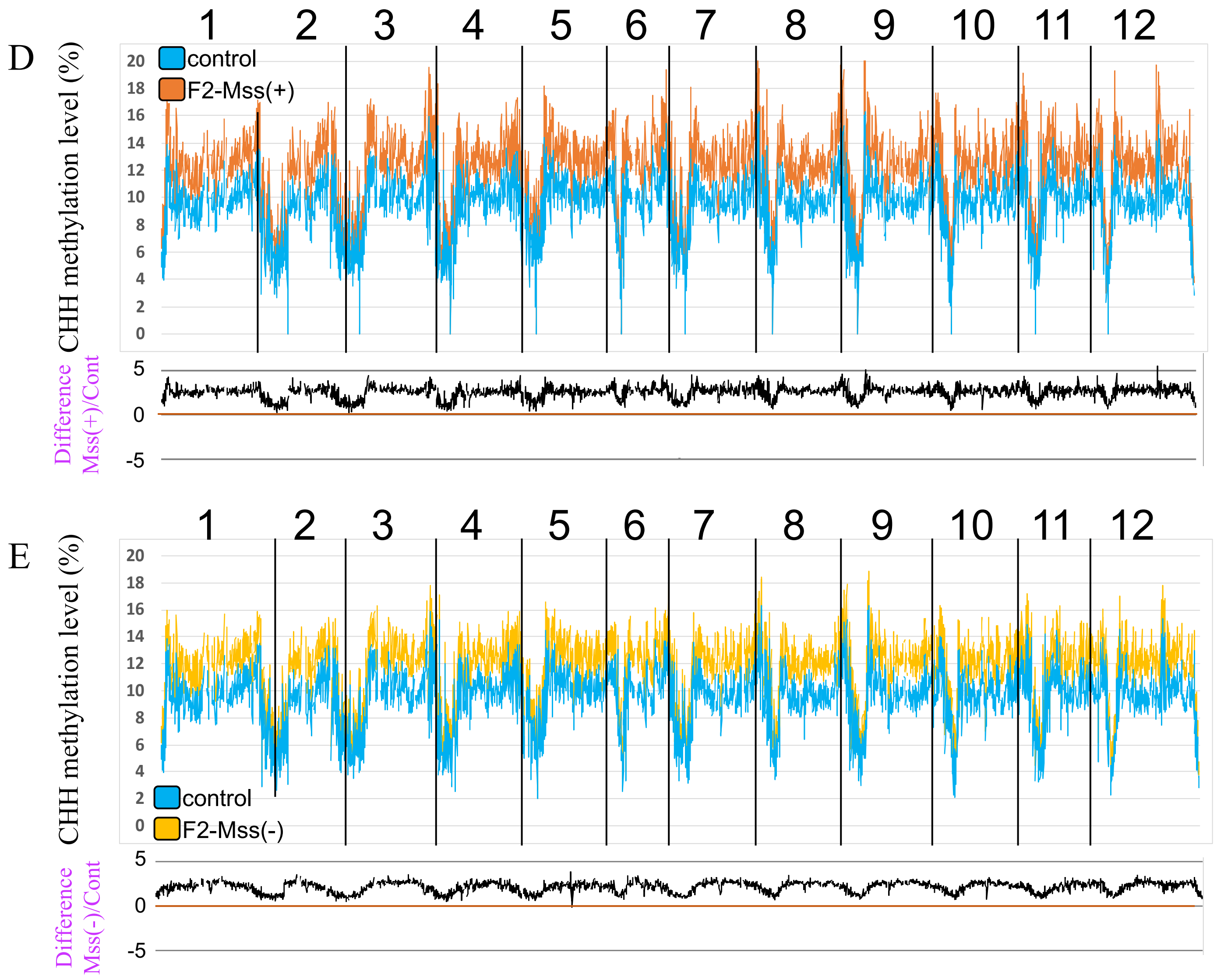


**Sup Figure 5.** Methylation along the 12 chromosomes of tomato, calculated from non-overlapping 200 kb bins. The average methylation levels were determined by combining the different plants for each genotype. Only cytosines covered by at least five reads were considered and only bins containing at least 10 valid cytosines were kept. Control: *pFIL::LhG4* driver line.

### Sup Figure 6

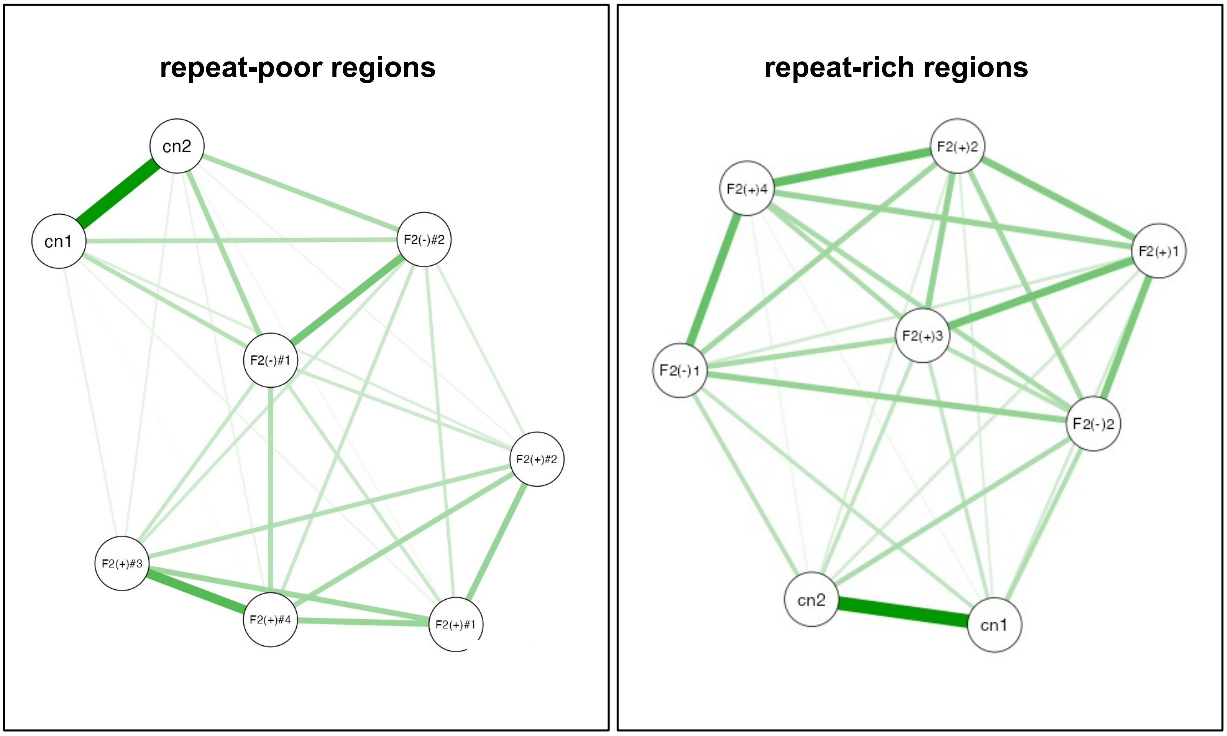


**Sup Figure 6.** Correlation network diagram constructed using Spearman's correlation coefficients to illustrate the relationships among the CG methylation bins depicted in the boxplots of Figure 3B. The lines represent the strength of the correlation: wider lines mean a stronger correlation. The spacing between the nodes is also indicative of the correlation level. Only correlations that are statistically significant (P-value < 0.05) are shown. cn, control driver line plants #1 and #2.

Sup Figure 7

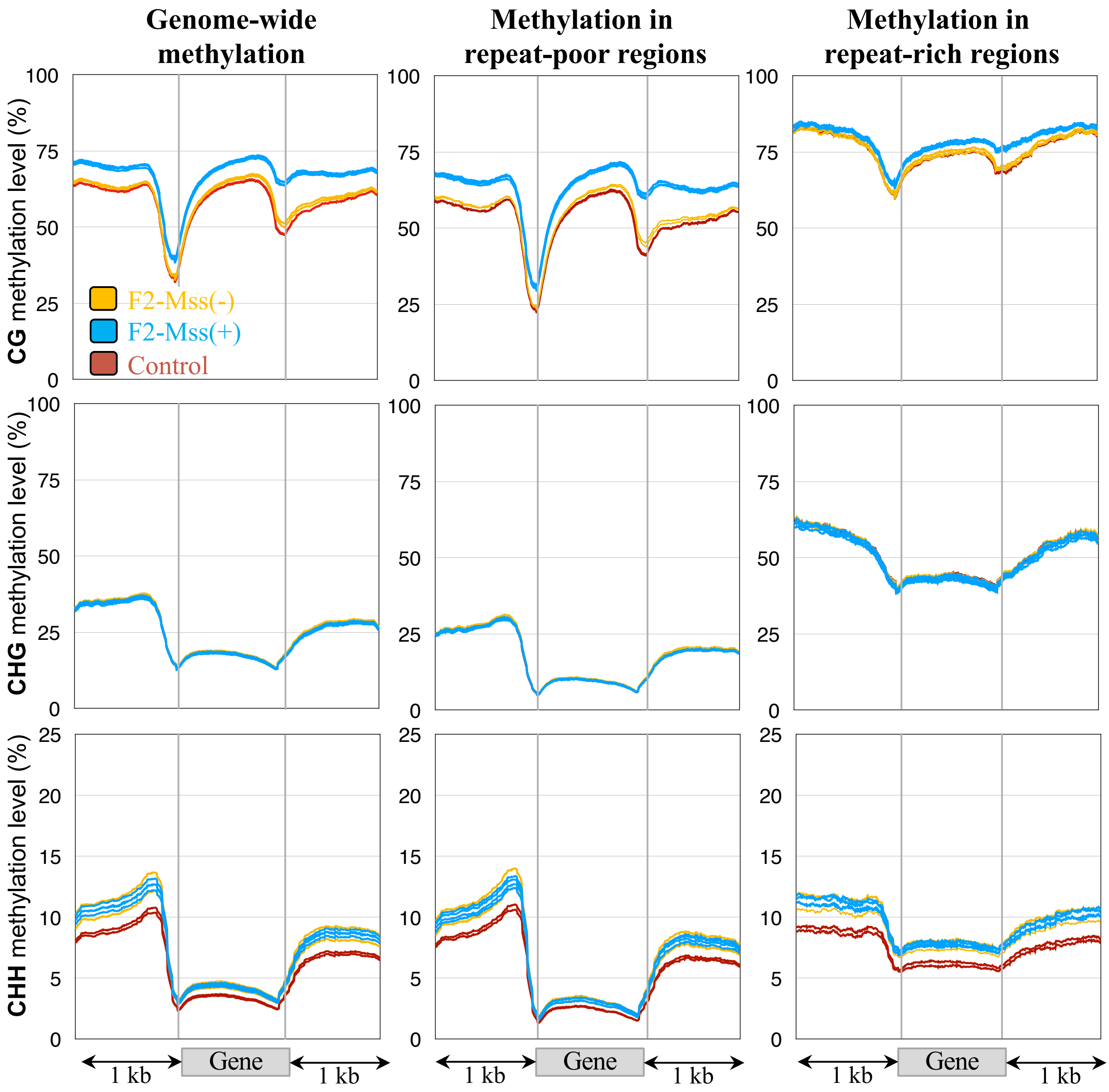


**Sup Figure 7.** Patterns of methylation in genes in the four Mss-F2(+) plants expressing *disM.SssI* (blue lines) and the control drive lines (orange lines). The average methylation level of genes was determined by dividing the corresponding annotated regions into 100 bp bins. Regions located 1 kb upstream and 1 kb downstream of genes are shown. The repeat-rich and repeat-poor regions were defined as previously described (Jouffroy et al., 2016), using the SL2.50 version of the genome assembly.

Sup Figure 8

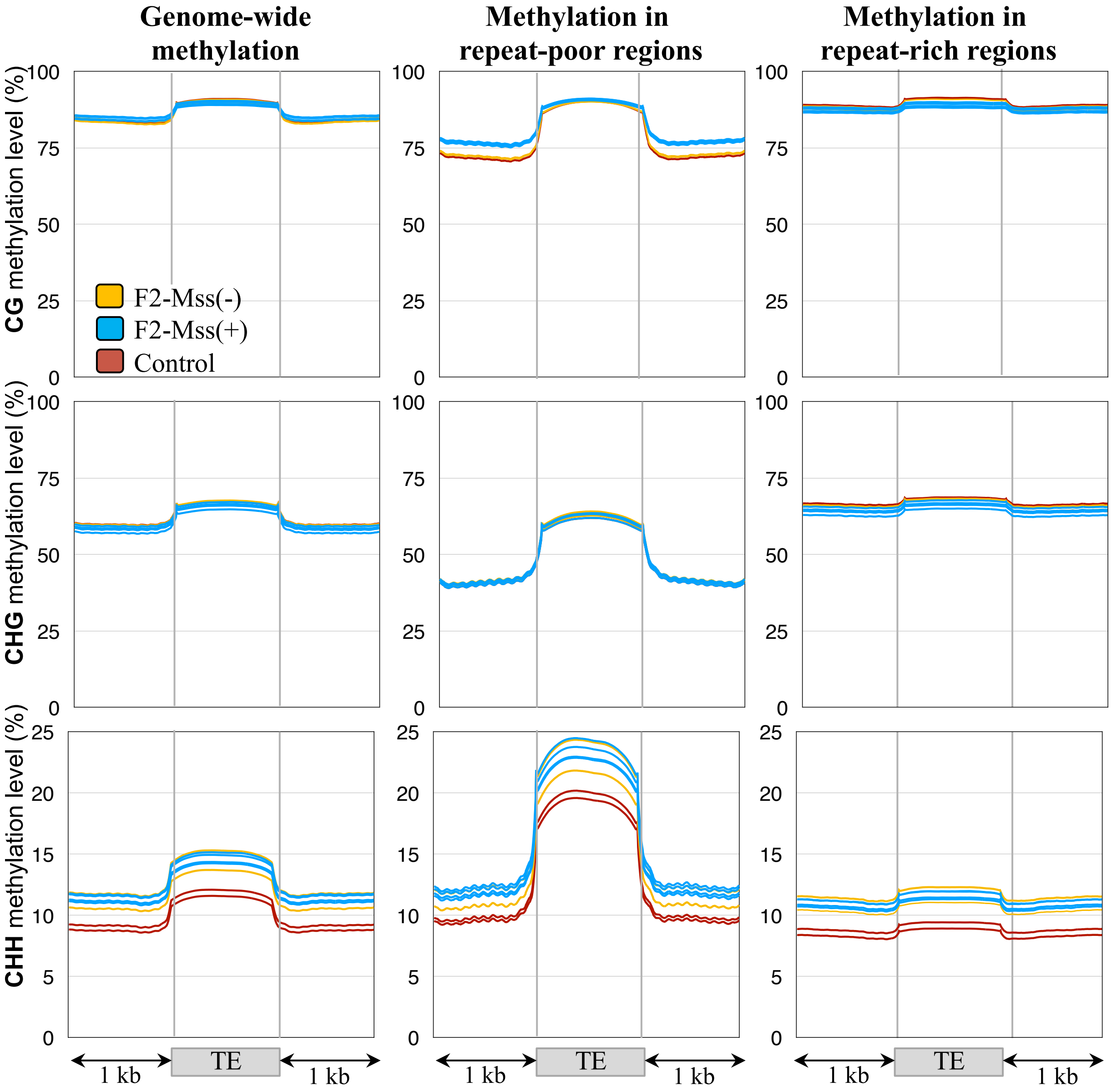


**Sup Figure 8.** Patterns of methylation in Transposable Elements (TEs) in the four Mss-F2(+) plants expressing *disM.SssI* (blue lines) and the control drive lines (orange lines). The average methylation level of TEs was determined by dividing the corresponding annotated regions into 100 bp bins. Regions located 1 kb upstream and 1 kb downstream of the TEs are shown. The repeat-rich and repeat-poor regions were defined as previously described (Jouffroy et al., 2016), using the SL2.50 version of the genome assembly.

Sup Figure 9

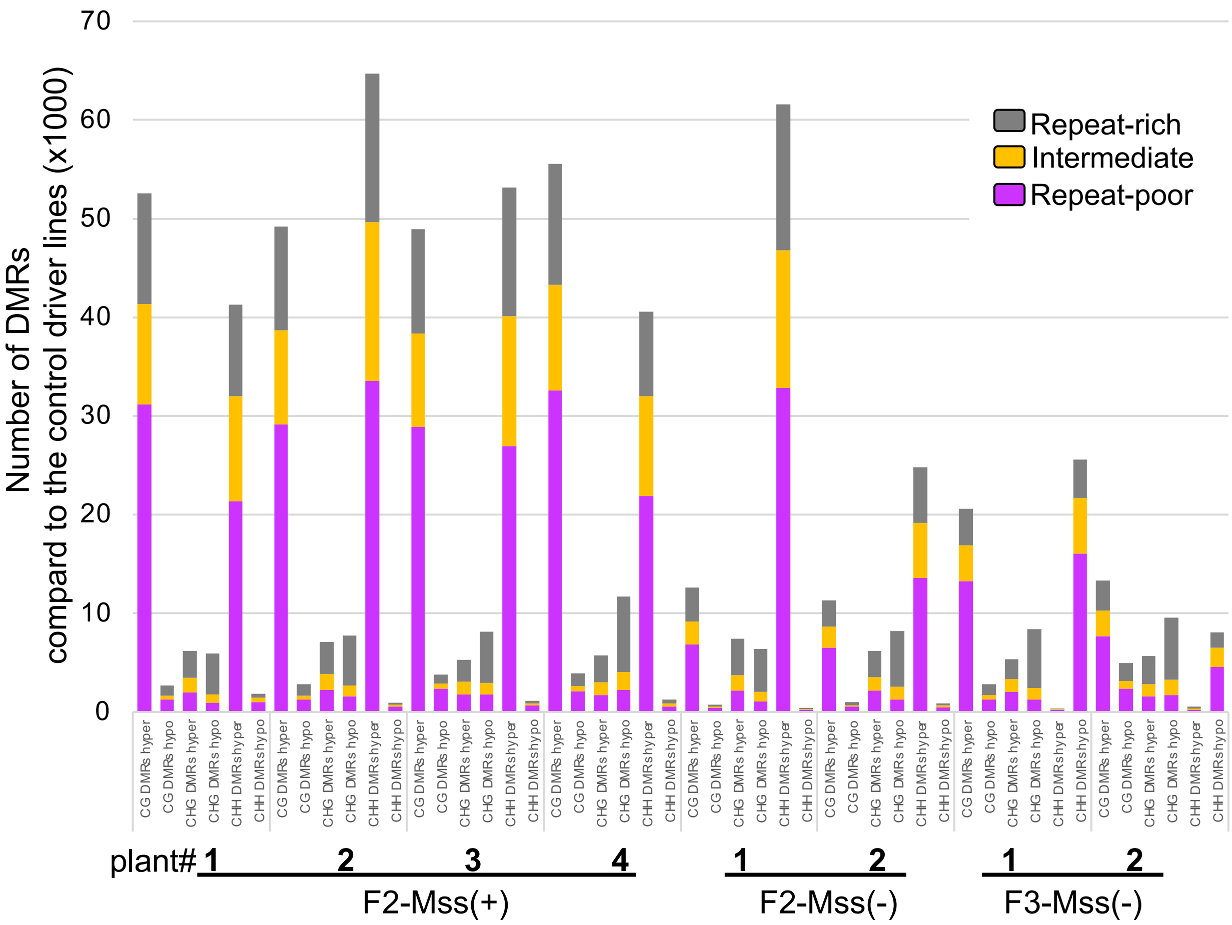


**Sup Figure 9.** Number of DMRs detected in F2 and F3 plants compared to the control driver line plants.

Repeat-Poor regions (RP), Repeat-Intermediate regions (INT), and Repeat-Rich regions (RR) were defined as previously described (Jouffroy et al., 2016), using the SL2.50 version of the genome assembly.

Sup Figure 10

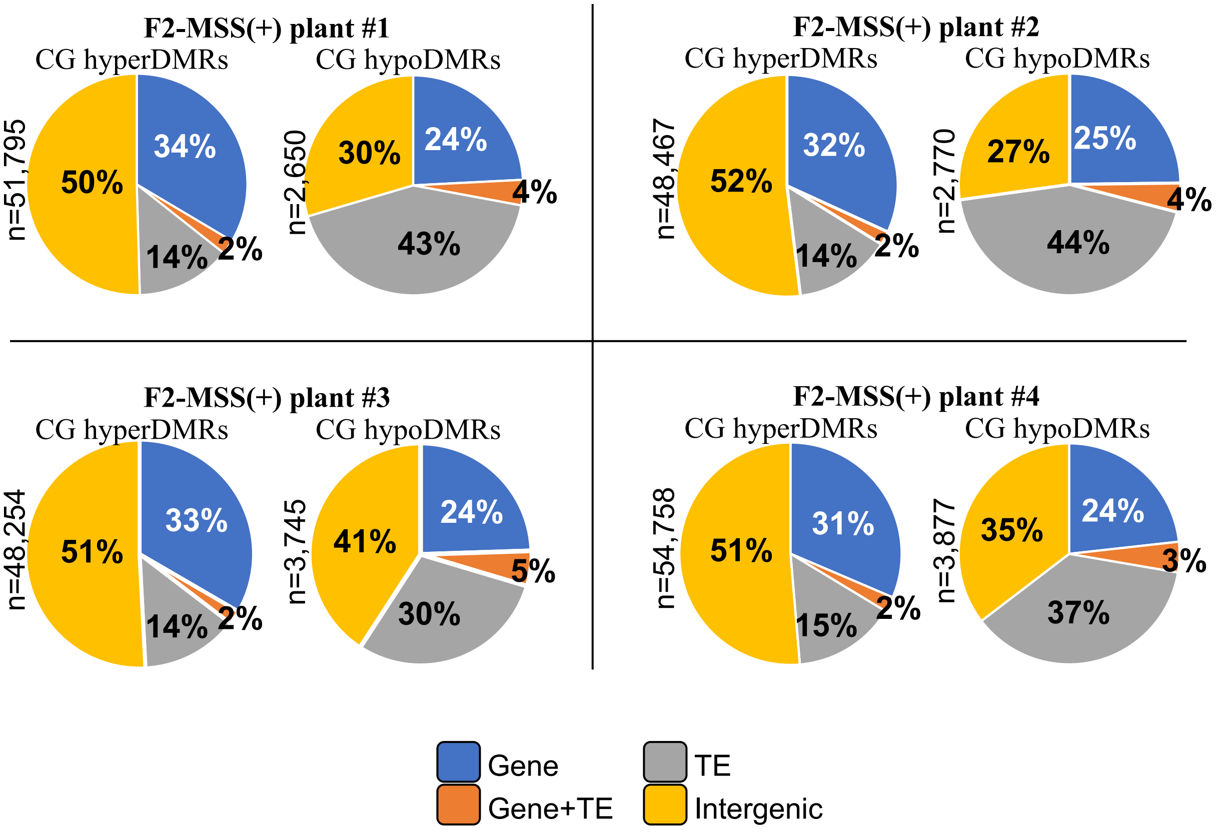


**Sup Figure 10.** Nature of the CG DMRs identified between the F2-Mss(+) plants and the driver line control plants. "Gene+TE" are DMRs overlapping with both genes and transposons, "Gene", DMRs overlapping with genes, and "TE" DMRs overlapping with Transposable Elements (TEs). All other DMRs were classified as "Intergenic".

Sup Figure 11

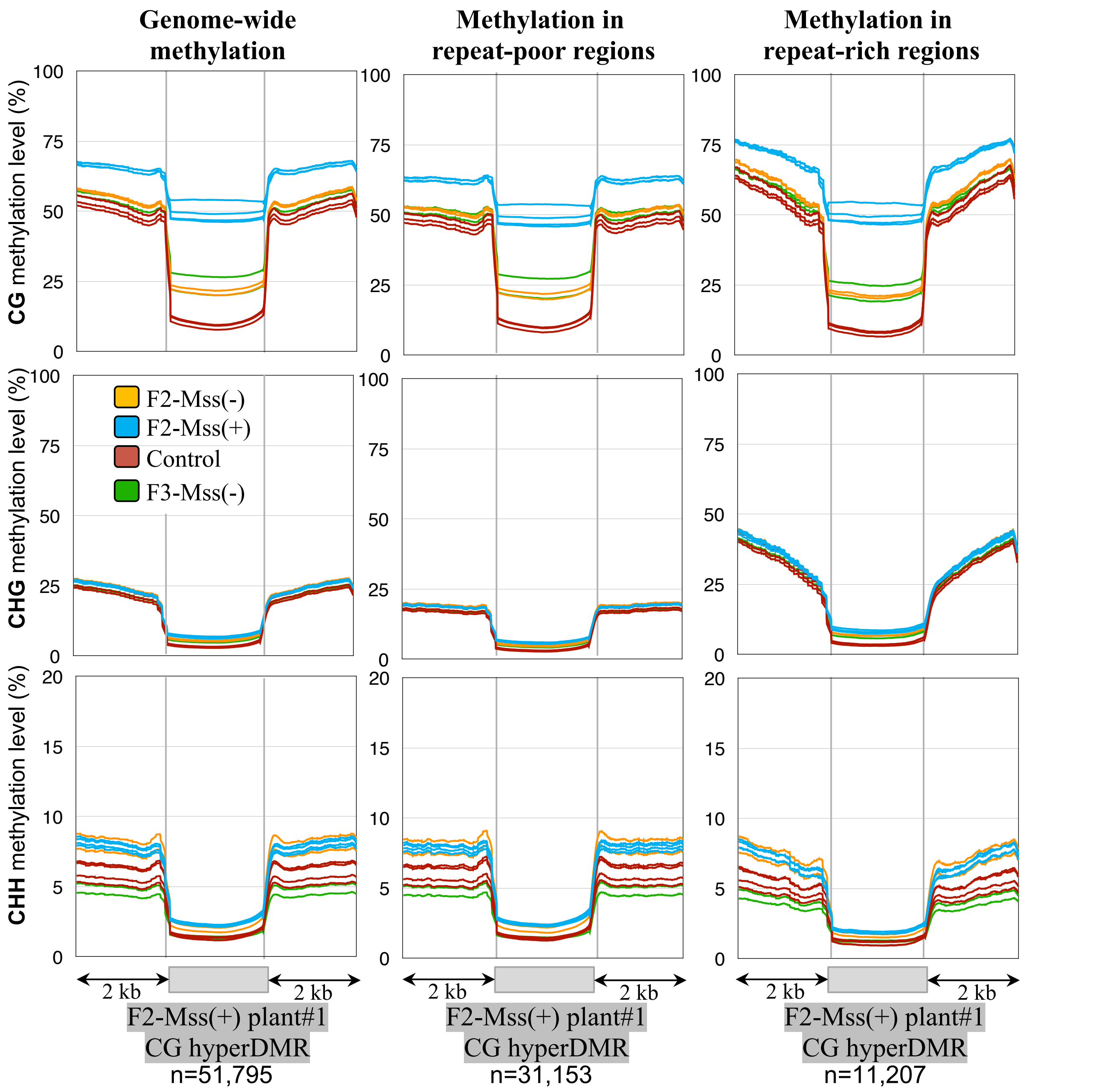


**Sup Figure 11.** Patterns of methylation for CG hyperDMRs identified between F2-Mss(+) plant#1 and driver line controls.

Metaprofiles showing the methylation levels in the F2-Mss(+) transgenic lines expressing both the *pFIL::LhG4* and *pOP::disM.SssI* transgenes (the four blue lines), the control driver line expressing only *pFIL::LhG4* (the four red lines), the F2-Mss(-) plants carrying no transgenes (the two orange lines) and the F3-Mss(-) plants carrying only the *pFIL:LhG4* transgene (the two green lines). The average methylation level of DMRs was determined by dividing the corresponding annotated regions into 100bp-bins. Regions located 2 kb upstream and 2 kb downstream of the CG hyperDMR are shown.

Sup Figure 12

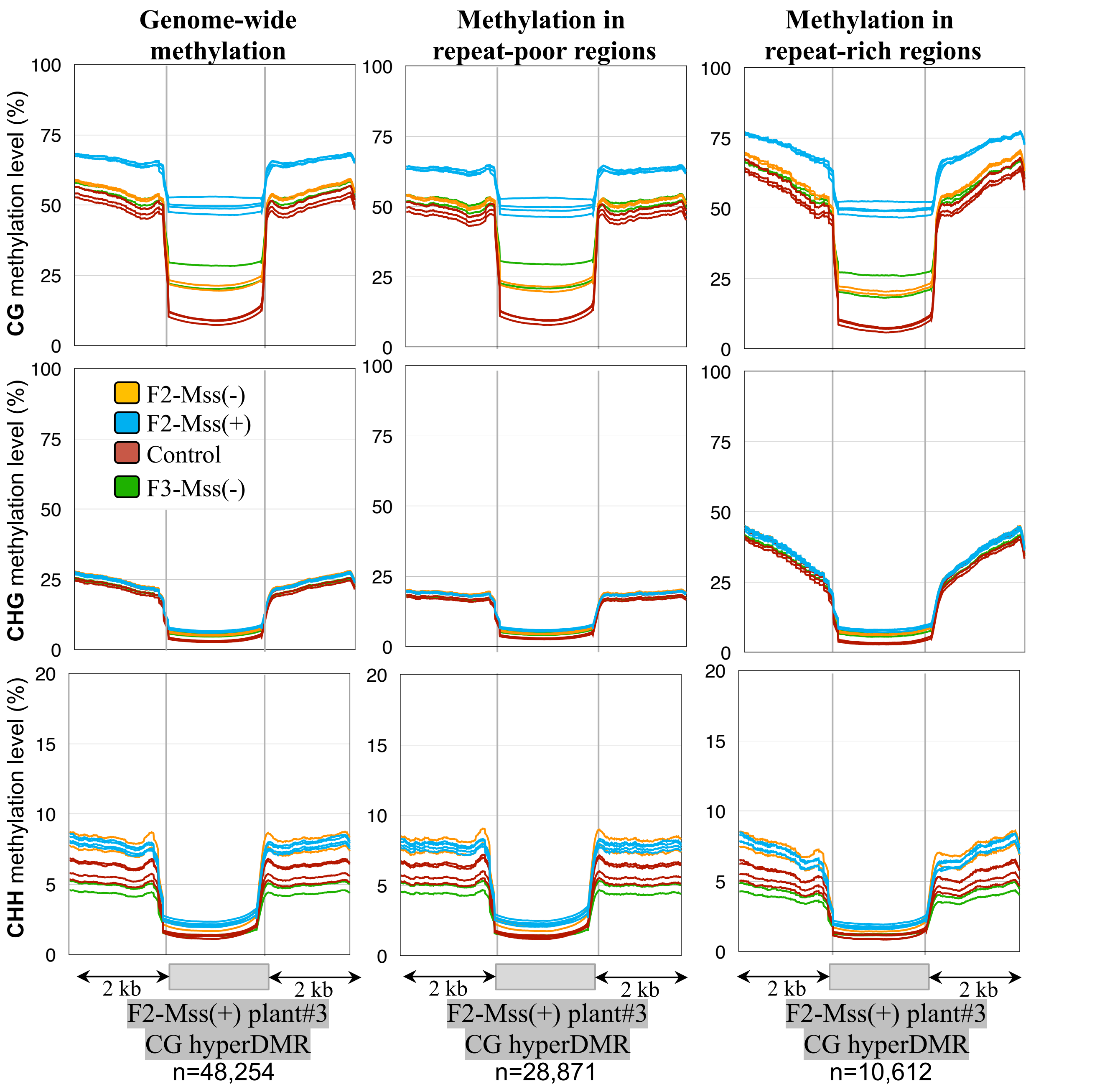


**Sup Figure 12.** Patterns of methylation for CG hyperDMRs identified between F2-Mss(+) plant#3 and driver line controls.

Metaprofiles showing the methylation levels in the F2-Mss(+) transgenic lines expressing both the *pFIL::LhG4* and *pOP::disM.SssI* transgenes (the four blue lines), the control driver line expressing only *pFIL::LhG4* (the four red lines), the F2-Mss(-) plants carrying no transgenes (the two orange lines) and the F3-Mss(-) plants carrying only the *pFIL:LhG4* transgene (the two green lines). The average methylation level of DMRs was determined by dividing the corresponding annotated regions into 100bp-bins. Regions located 2 kb upstream and 2 kb downstream of the CG hyperDMR are shown.

Sup Figure 13

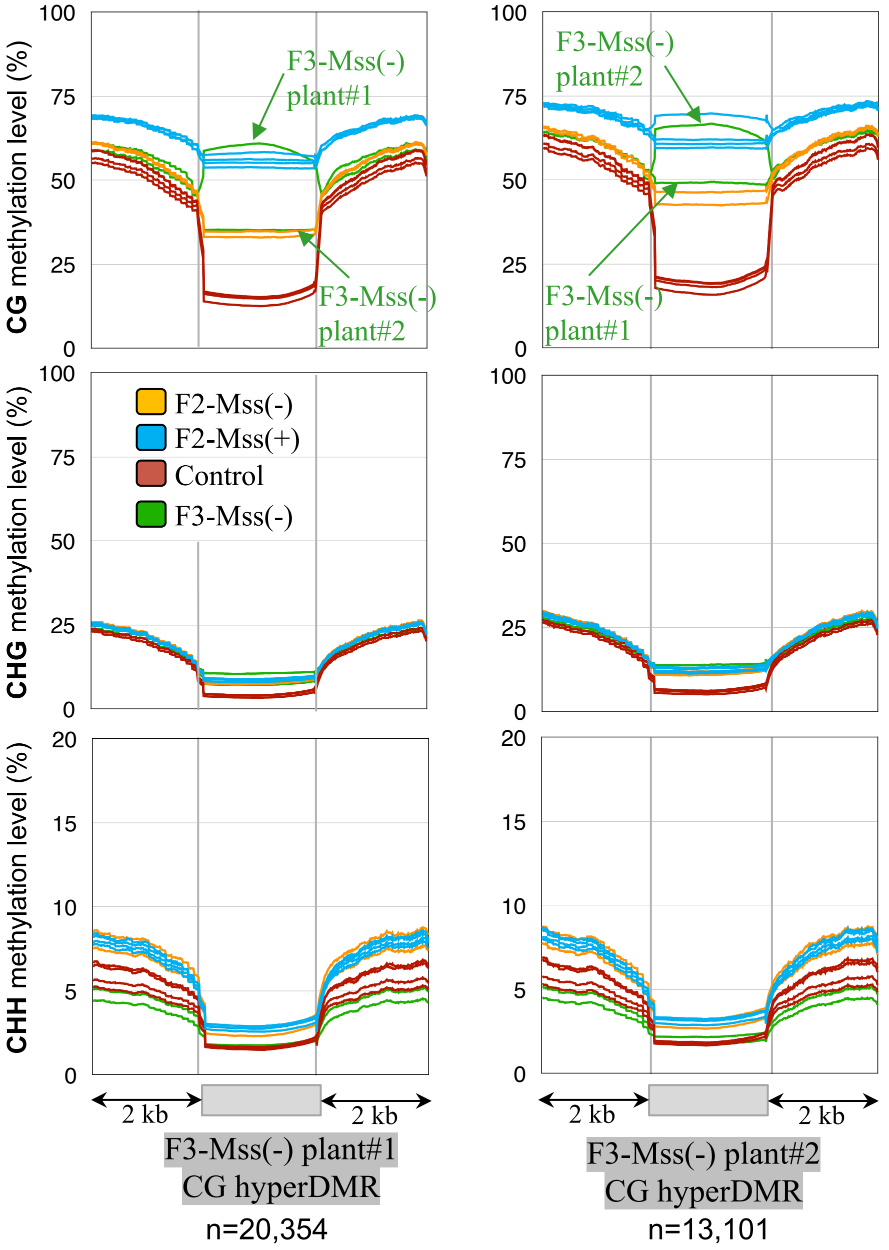


**Sup Figure 13.** Patterns of methylation for CG hyperDMRs identified between F3-Mss(-) and driver line control plants.

Metaprofiles show the methylation levels in the four F2-Mss(+) transgenic lines expressing both *pFIL::LhG4* and *pOP::disM.SssI* (blue lines), the control driver lines expressing only *pFIL:LhG4* (red lines), the F2-Mss(-) plants carrying no transgenes (orange lines) and the F3-Mss(-) plants carrying only the *pFIL::LhG4* transgene (green lines). The average methylation level of DMRs was determined by dividing the corresponding annotated regions into 100bp-bins. Regions located 2 kb upstream and 2 kb downstream of the CG hyperDMR are shown.

Sup Figure 14

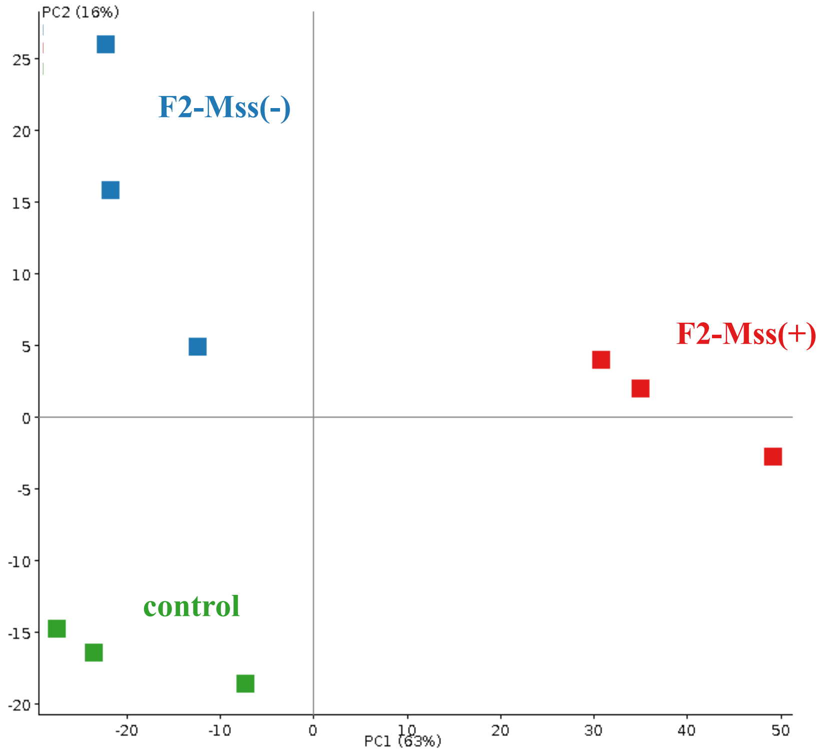


**Sup Fig 14.** Principal Component Analysis (PCA) plots of RNAseq data

Sup Figure 15

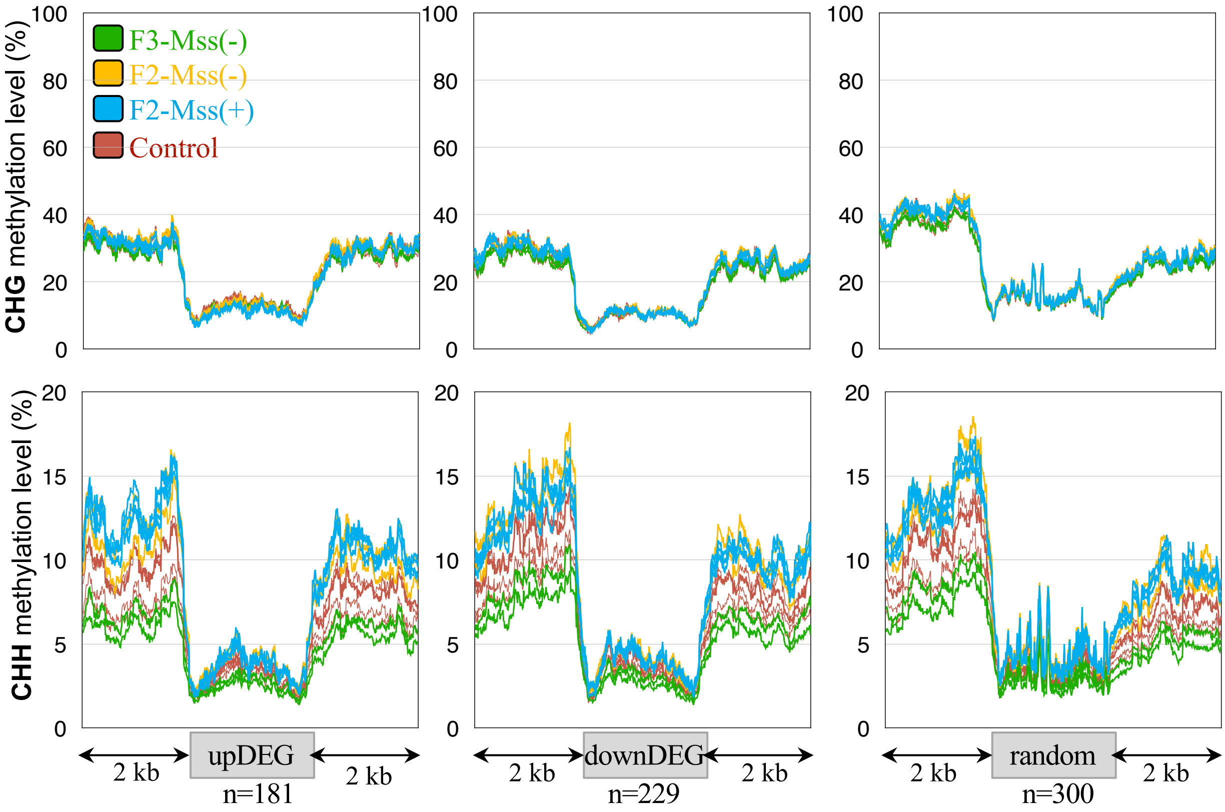


**Sup Fig 15.** Methylation levels of genes that are up- (upDEG) or downregulated (downDEG) (> 1 ou < -1 log2FC) between the control and plants F2-Mss(+) expressing the methylase.

The average methylation levels of Differentially Expressed Genes (DEGs) were determined by dividing the genes into 100-bp bins. Regions located 2 kb upstream and 2 kb downstream are shown.

Random : set of 300 genes chosen randomly.

### Sup Figure 16

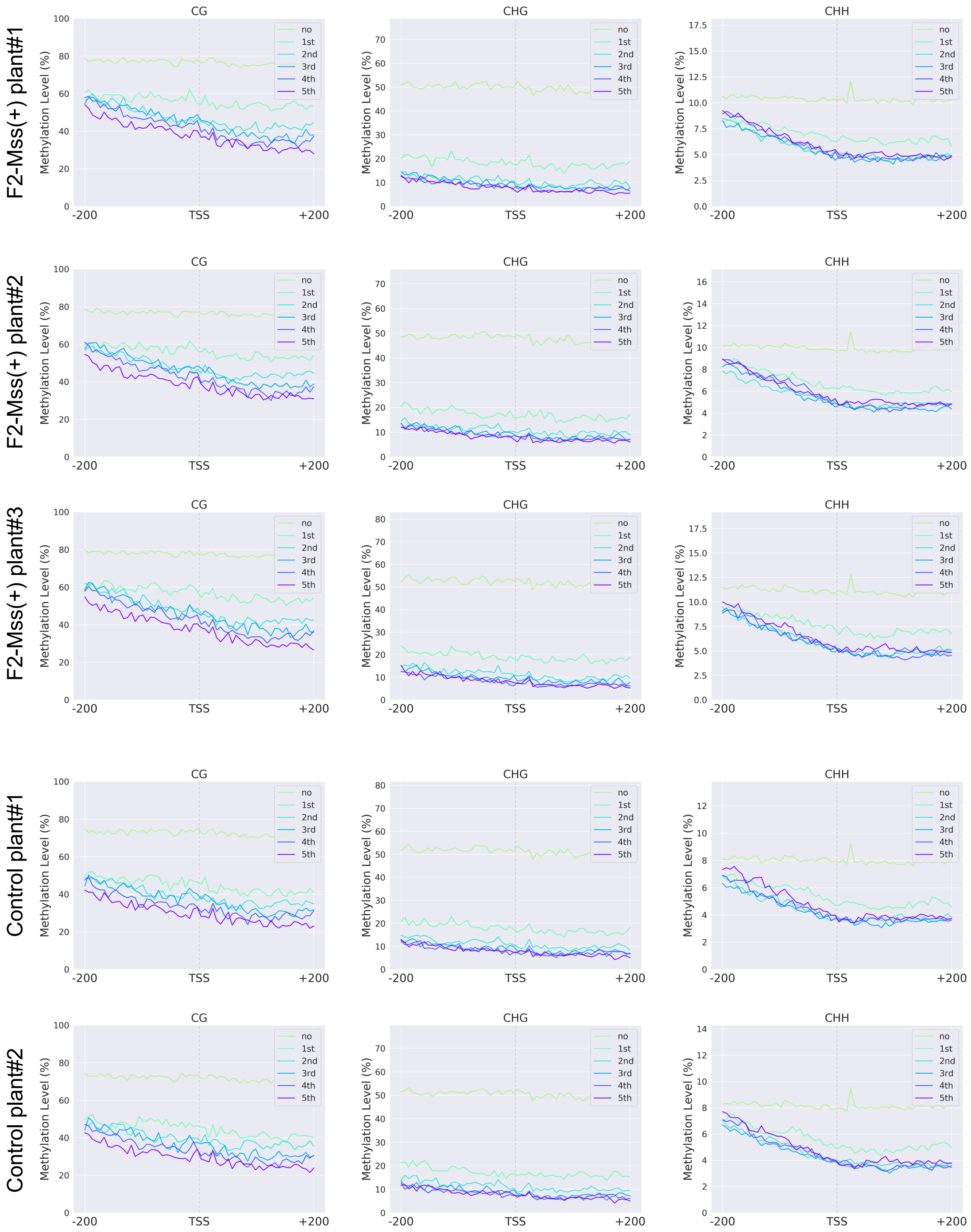


**Sup Fig 16.** Average methylation level profiling according to different expression groups around the TSS (+/- 200 bp) of F2-Mss(+) and control lines. Genes are grouped as non-expressed genes and 5 quantiles of expressed genes according to the gene expression level groups from low to high; the 1st quintile is the lowest, and the 5th is the highest.
